# Multisensory pathogen detection drives rapid escape and shapes host–microbe interactions in *Drosophila* larvae

**DOI:** 10.64898/2025.12.18.695079

**Authors:** Olivier Zugasti, Julien Royet

**Author notes:** Authors contributed equally.

## Abstract

Larvae of many insects develop immersed in decomposing substrates densely populated with microbes, yet how they detect and respond to pathogenic threats during early life remains poorly understood. Here, we show that *Drosophila melanogaster* larvae exhibit a previously unrecognized rapid escape behavior triggered by food contaminated with metabolically active *Erwinia carotovora carotovora 15* (*E. cc^15^*), a natural bacterial pathogen of flies and plants. Stationary-phase cells fail to elicit avoidance, demonstrating that larval detection of *E. cc^15^* depends on bacterial metabolic activity rather than on its mere presence. Using targeted genetic manipulations, we identify two chemosensory pathways required for this response: a gustatory input mediated by the aversion receptor Gr33a and an olfactory input mediated by the Or49a–Orco complex. Disrupting either pathway abolishes escape, revealing that larvae integrate complementary contact-dependent and volatile cues to evaluate microbial danger. Functionally, escape enables larvae to reach uncontaminated food and partially mitigates the developmental impact of early pathogen exposure. However, dispersing larvae also transfer viable bacteria to new substrates, indicating that this defensive response can simultaneously promote pathogen spread. Together, these findings establish the first example of a rapid, multisensory escape behavior induced by a natural pathogen in *Drosophila* larvae and provide a tractable model for dissecting how microbial cues guide defensive decision-making and influence pathogen dissemination.

## INTRODUCTION

Animals developing in microbe-rich environments must continuously balance the need to acquire nutrients with the risk of encountering pathogenic microorganisms [1,2]. In holometabolous insects such as *Drosophila melanogaster*, this constraint is particularly pronounced during the larval stage, when individuals feed almost continuously within decaying fruits and fermenting substrates densely populated with bacteria and yeasts [3,4]. This ecology has shaped a robust and well-characterized innate immune system in both larvae and adults, enabling the detection and control of diverse microbial threats [5]. Several natural pathogens of *Drosophila*, including *Erwinia carotovora carotovora* 15 (*E. cc^15^*), *Pseudomonas entomophila*, and *Serratia marcescens*, have therefore become powerful models for dissecting host–microbe interactions and infection-induced physiological alterations [6–8].

Among these microbes, *E. cc^15^* is a Gram-negative phytopathogen of the *Enterobacteriaceae* family that proliferates in damaged plant tissues and causes soft-rot disease [9]. Its ecological niche overlaps with that of *Drosophila*, and adult flies can naturally acquire and disseminate *Erwinia* between plant hosts [10–12]. Ingestion of *E. cc^15^* triggers strong immune activation, including systemic immune deficiency pathway (IMD) stimulation through detection of diaminopimelate-type peptidoglycan (PGN) [6,13–15] and, in adults, uracil-dependent DUOX activation in the gut epithelium, leading to the production of reactive oxygen species [16]. *E. cc^15^*also expresses the virulence factor Evf (*Erwinia* virulence factor), which promotes bacterial persistence in the larval gut and modifies host physiology, including delaying gut clearance and altering feeding behavior [17,18]. These features indicate that *Drosophila* naturally encounter *E. cc^15^*and raise the possibility that the host may have evolved strategies to detect and limit bacterial contact before infection becomes established.

Behavioral defenses such as avoidance, withdrawal, and refuge seeking constitute a first line of protection that reduces pathogen exposure before immune mechanisms are engaged across animal taxa [19]. In adult *Drosophila*, several dedicated sensory pathways mediate avoidance of harmful microbes or their metabolites, including olfactory detection of fungal geosmin and phenolic volatiles through the odorant receptors Or56a [20] and Or46a [21], and gustatory detection of bacterial lipopolysaccharides (LPS) via the TRPA1 nociceptor in *Gr66a*-expressing bitter neurons [22]. These findings demonstrate that adults rely on both olfaction and gustation to evaluate microbial risk.

In contrast, much less is known about corresponding strategies in larvae, whose sensory architecture, foraging habits, and ecological constraints differ substantially [23–25]. This gap is notable given that larvae develop immersed in their food substrate, ingesting large quantities of microbes while relying primarily on close-range chemical cues to assess food quality. Whether larvae can detect natural pathogens, whether they mount behavioral responses to limit exposure, and which sensory pathways mediate such detection all remain unresolved. Moreover, if such behaviors exist, their consequences for larval survival and their potential role in pathogen dissemination have not been explored.

Here, we address these questions by analyzing larval behavioral responses to food contaminated with *E. cc^15^* and by dissecting the sensory circuits that mediate pathogen-induced escape and its ecological consequences.

## RESULTS

### Metabolically active *E. cc^15^* Triggers dispersal behavior in *Drosophila* larvae

To assess larval behavior when exposed to a source of food contaminated with *E. cc^15^*, baker’s yeast (*Saccharomyces cerevisiae*), a nutritive substrate supporting optimal larval growth [26], was mixed with increasing concentrations of stationary-phase *E. cc^15^* cultured overnight (S1A Fig). This yeast–bacteria mixture was then placed on nutrient-rich LB agar plates and left to incubate at room temperature for 30 minutes, a duration that promotes bacterial regrowth in static liquid culture (S1B Fig). Wild-type (w¹¹¹⁸) second-instar larvae were subsequently transferred directly onto the contaminated food for behavioral assays (Fig 1A).

**Fig 1.**
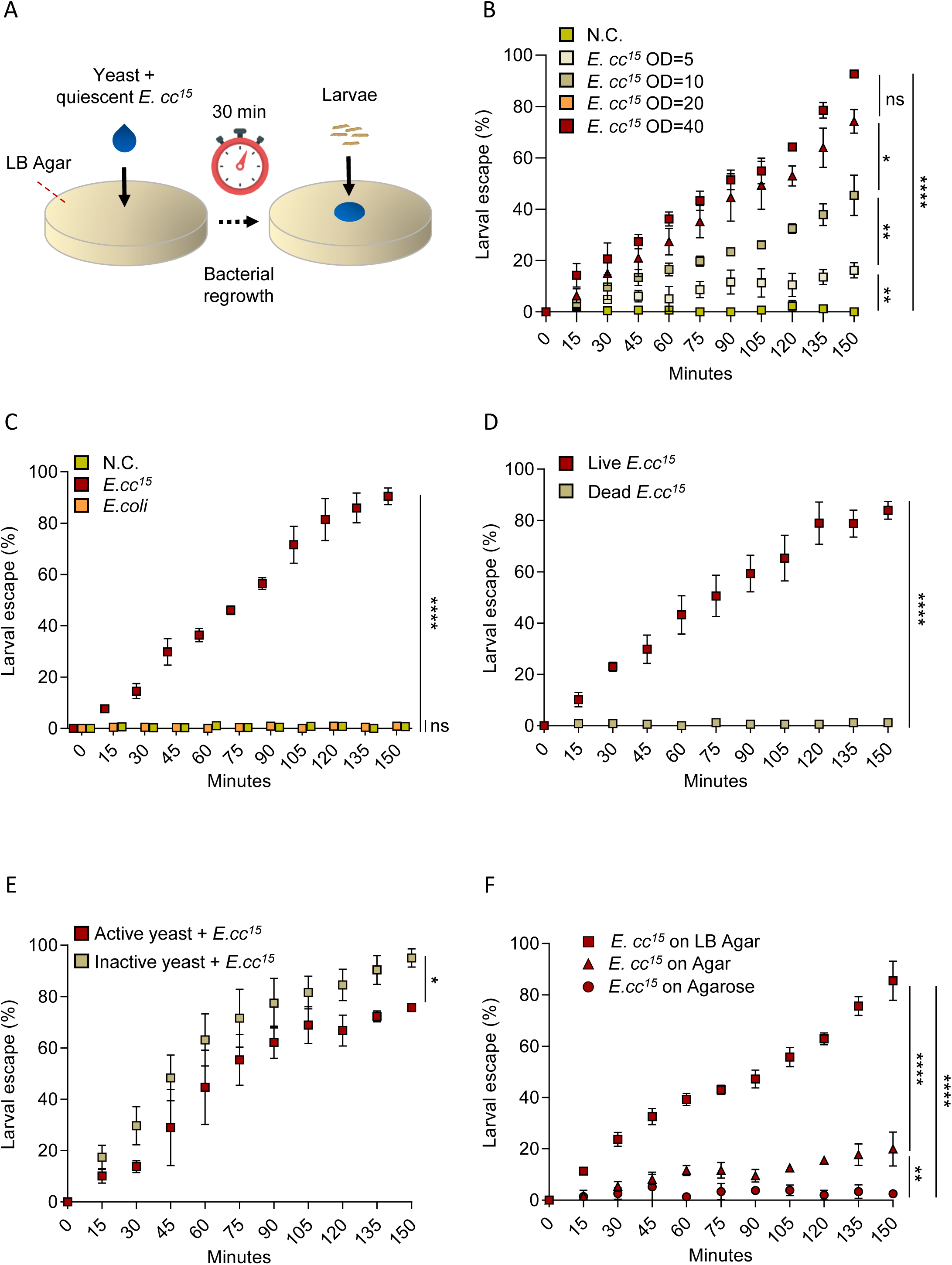
Metabolically active *E. cc^15^* cells trigger rapid and dose-dependent escape of *Drosophila* larvae. **(A)** Schematic of the assay used to evaluate larval responses to *E. cc^15^*-contaminated food. **(B)** Quantification of larval dispersal from uncontaminated yeast (N.C.) and yeast contaminated with increasing concentrations of *E. cc^15^*(OD_600_ = 5, 10, 20, and 40) over time. **(C)** Comparison of larval dispersal on uncontaminated yeast versus yeast contaminated with *E. cc^15^* or *E. coli* at identical bacterial densities (OD_600_ 40). **(D)** Dispersal of larvae exposed to live versus heat-killed *E. cc^15^*. **(E)** Larval behavior on mixtures of yeast and *E. cc^15^* prepared with metabolically active or inactivated yeast. **(F)** Effect of nutrient-rich LB agar, nutrient-poor plain agar, and nutrient-free agarose substrates on *E. cc^15^*-induced escape. For this and all subsequent figures, quantified dispersal assay results are shown as mean ± SD of technical triplicates from a representative biological experiment. Each experiment was independently repeated at least three times, using groups of ≥40 larvae per replicate. Statistical analyses were performed using a mixed-effects model (REML) with time and condition as fixed factors, (ns = not significant; *p < 0.05; **p < 0.01; ***p < 0.001; ****p < 0.0001). Comparisons between selected conditions are shown. For all experiments, raw data and full statistical outputs are provided in S1 Data.

Exposure to *E. cc^15^*-contaminated food caused larvae to disperse within minutes, and this response grew stronger both over time and with higher bacterial concentrations (Fig 1B). At bacterial densities equivalent to OD_600_ 20–40, more than 80% of the larvae had dispersed within 2.5 hours. Lower concentrations (OD_600_ 5–10) produced a weaker initial reaction. Once larvae moved away from the contaminated food, they remained off the substrate for the rest of the observation period, suggesting they actively avoided returning to it (S2A Fig). After 24 hours, nearly all larvae had abandoned the *E. cc^15^*-contaminated yeast at every concentration tested, whereas larvae consistently stayed on uncontaminated food (S2A Fig). Thus, the strength of the dispersal response scaled with the bacterial load. Based on the rapidity and reproducibility of the response at OD_600_ 40, this bacterial concentration was selected for subsequent experiments unless otherwise indicated.

To evaluate the specificity of this behavioral response, larvae were exposed to yeast mixed with *Escherichia coli* (*E. coli*), a non-pathogenic member of the *Enterobacteriaceae* family, prepared from stationary-phase cultures and applied to LB agar plates using the same procedure as for *E. cc^15^* (S1C–D Fig). While larvae rapidly abandoned food contaminated with *E. cc^15^*, they remained on yeast containing *E. coli* as well as on uncontaminated yeast (Fig 1C, S3 Fig, S1-3 Movies). This indicates that the dispersal behavior is triggered by cues specific to *E. cc^15^* rather than by the mere presence of bacteria.

The robustness of this dispersal response was further verified across multiple *Drosophila melanogaster* genetic backgrounds including *yellow white*, *Oregon-R*, and *Canton-S* larvae, all of which displayed rapid escape from *E. cc^15^*-contaminated food (S4 Fig). To evaluate the temporal stability of this behavior, yeast– *E. cc^15^* mixtures stored on LB agar plates for 2, 4, or 6 days were tested. In every case, larvae continued to disperse robustly and extensively, demonstrating that this avoidance behavior is stable and independent of the age of the bacterial preparation (S5 Fig).

Additional experiments tested whether larval dispersal depends on bacterial viability and metabolic activity. Only live *E. cc^15^* bacteria induced dispersal; heat-killed bacteria produced no response (Fig 1D). Moreover, larvae dispersed equally from food containing live *E. cc^15^* mixed with either metabolically active or inactive yeast (Fig 1E), demonstrating that the behavior is driven specifically by bacterial cues rather than by any yeast-derived signals.

To further assess the role of bacterial metabolism, yeast– *E. cc^15^* mixtures were placed on nutrient-rich LB agar, nutrient-poor simple agar, or nutrient-free agarose. Larvae dispersed rapidly on LB agar, showed a weaker response on plain agar, and failed to disperse on agarose, where bacterial activity is expected to be minimal (Fig 1F and S6 Fig). These findings show that larval escape behavior requires metabolically active bacteria, whose activity is shaped by the nutrient content of the substrate.

Taken together, these results demonstrate that *Drosophila melanogaster* larvae exhibit a rapid, strong, and specific dispersal response to *E. cc^15^*-contaminated food. This avoidance behavior is triggered by signals from nutrient-responsive bacteria rather than by bacterial presence alone, and it depends on substrate-driven bacterial metabolic activity as a key component of this contact-dependent response.

### Larval avoidance of *E. cc^15^* emerges after initial non-selective food approach

To determine whether *Drosophila* larvae require direct contact with bacteria to initiate dispersal, an avoidance assay was performed (Fig 2A). Larvae were positioned at a distance from a food source under three conditions: uncontaminated yeast, yeast inoculated with *E. coli*, or yeast inoculated with *E. cc^15^*. In all conditions, the larvae rapidly oriented toward the food and entered it, confirming an attraction to nutritional cues (Fig 2B). However, upon contacting yeast contaminated with *E. cc^15^*, larvae displayed a rapid escape response and left the substrate shortly after arrival (Fig 2B). By contrast, larvae that reached uncontaminated yeast or yeast containing *E. coli* remained on the food throughout the assay (Fig 2B). These observations indicate that larvae do not detect *E. cc^15^* at a distance but instead respond to cues perceived upon direct contact with the contaminated food.

**Fig 2.**
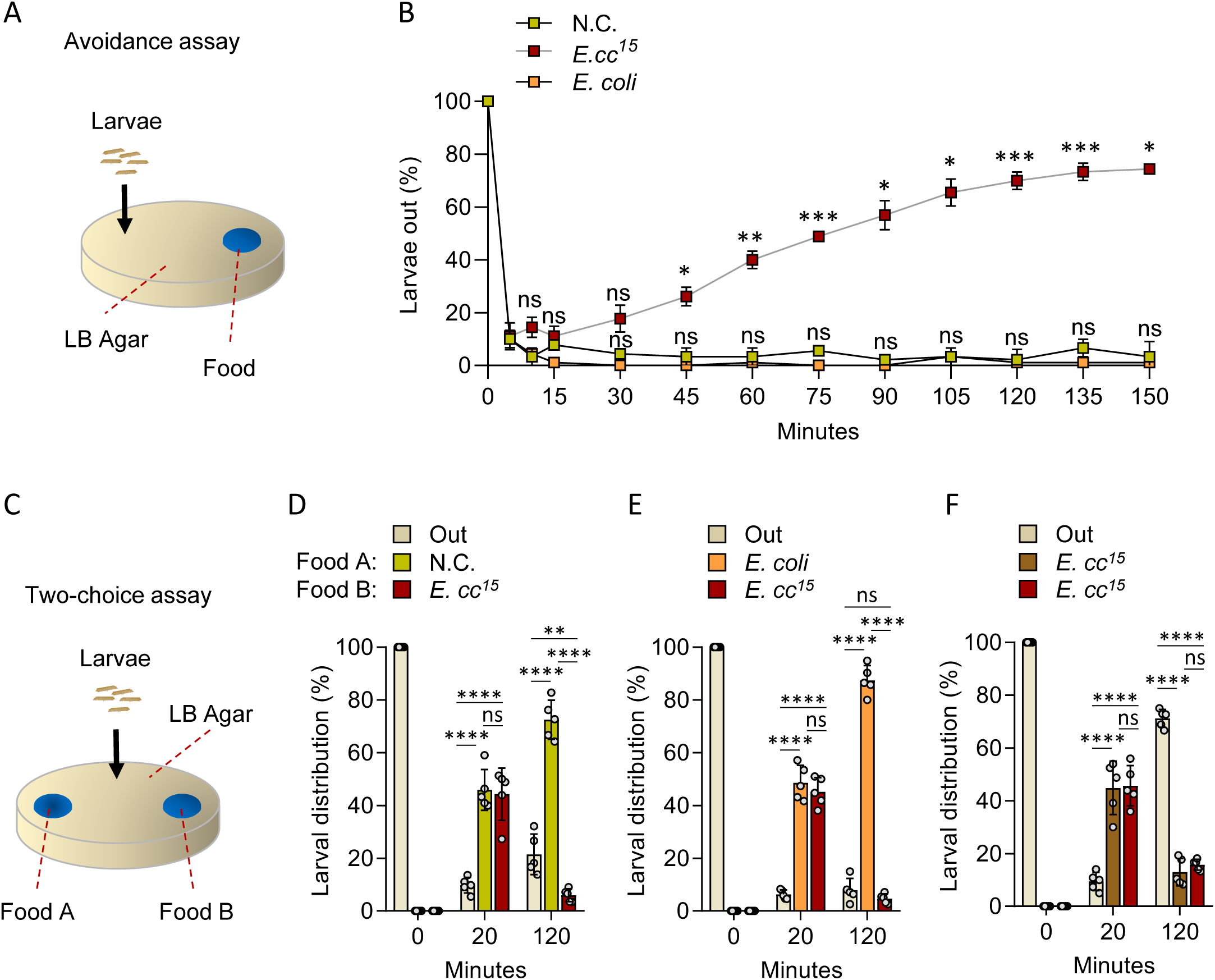
Initial non-selective approach precedes larval escape from *E. cc^15^*-contaminated food. **(A)** Schematic of the assay monitoring larval behavior toward a distant food source. **(B)** Quantification of larvae located outside the food over time when placed at a distance from uncontaminated yeast (N.C.) or yeast contaminated with *E. coli* or *E. cc^15^*. Data are presented as means ± SD of technical triplicates using groups of 30 Larvae. Statistical analyses were performed using a mixed-effects model (REML) with time and condition as fixed factors, followed by Tukey’s post-hoc tests comparing uncontaminated food with either *E. cc^15^*-contaminated or *E. coli*-contaminated food conditions (ns = not significant; *p < 0.05; **p < 0.01; ***p < 0.001; **p < 0.0001). **(C)** Schematic of the binary food-choice assay. **(D-F)** Distribution of larvae at 0, 20, and 120 min in food-choice assays when presented with paired patches: uncontaminated versus *E. cc^15^*-contaminated yeast (D); *E. coli* versus *E. cc^15^*-contaminated yeast (E); two *E. cc^15^*-contaminated patches (F). Data are presented as means ± SD. Individual points represent experimental replicates (each consisting of groups of ≥ *30* larvae). Statistical analyses were performed using Fisher’s exact test (ns = not significant; *p < 0.05; **p < 0.01; ***p < 0.001; ****p < 0.0001).

To examine how larvae adjust their behavior when presented with multiple food options, a two-choice assay was conducted (Fig 2C). Larvae were placed equidistant between two food patches, and three pairwise comparisons were tested: uncontaminated yeast versus yeast inoculated with *E. cc^15^*; *E. coli* versus *E. cc^15^*contaminated yeast; and two patches both containing *E. cc^15^*. Larval positions were recorded at 20 and 120 minutes to capture their initial orientation and later decision-making. After 20 minutes, larvae reached one of the food patches in all conditions without showing any preference (Fig 2D–F), indicating that early orientation is not influenced by bacterial contamination. By 120 minutes, however, clear differences emerged: larvae moved away from *E. cc^15^*-contaminated food and preferentially occupied uncontaminated or *E. coli*-associated food (Fig. 2D–E). When both patches contained *E. cc^15^*, larvae abandoned the food entirely and accumulated on the surrounding agar (Fig 2F), consistent with a broad avoidance response in the absence of an alternative food source. These findings demonstrate that *Drosophila* larvae initially approach food without discrimination but retreat upon contact with *E. cc^15^*-contaminated substrates, subsequently redirecting their movement toward innocuous food sources. Combined with previous evidence that dispersal depends on bacterial metabolic activity rather than mere presence, these results suggest that contact-based detection of *E. cc^15^* involves specific bacterial metabolites or virulence-associated factors.

### Major *E. cc^15^*-derived factors or canonical immune and nociceptive signaling do not mediate larval dispersal behavior

Metabolically active *E. cc^15^* produces multiple factors, including virulence proteins, metabolites, and structural components, that influence *Drosophila* physiology and potentially cue larval dispersal. The secreted virulence factor Evf, which promotes gut colonization and trigger systemic immune activation [17,27], was first assessed. Larvae exposed to yeast contaminated with either wild-type or *evf*-deficient *E. cc^15^* (*E. cc^15^ Δ evf*) [17] dispersed equally, indicating that Evf is not required (Fig 3A).

**Fig 3.**
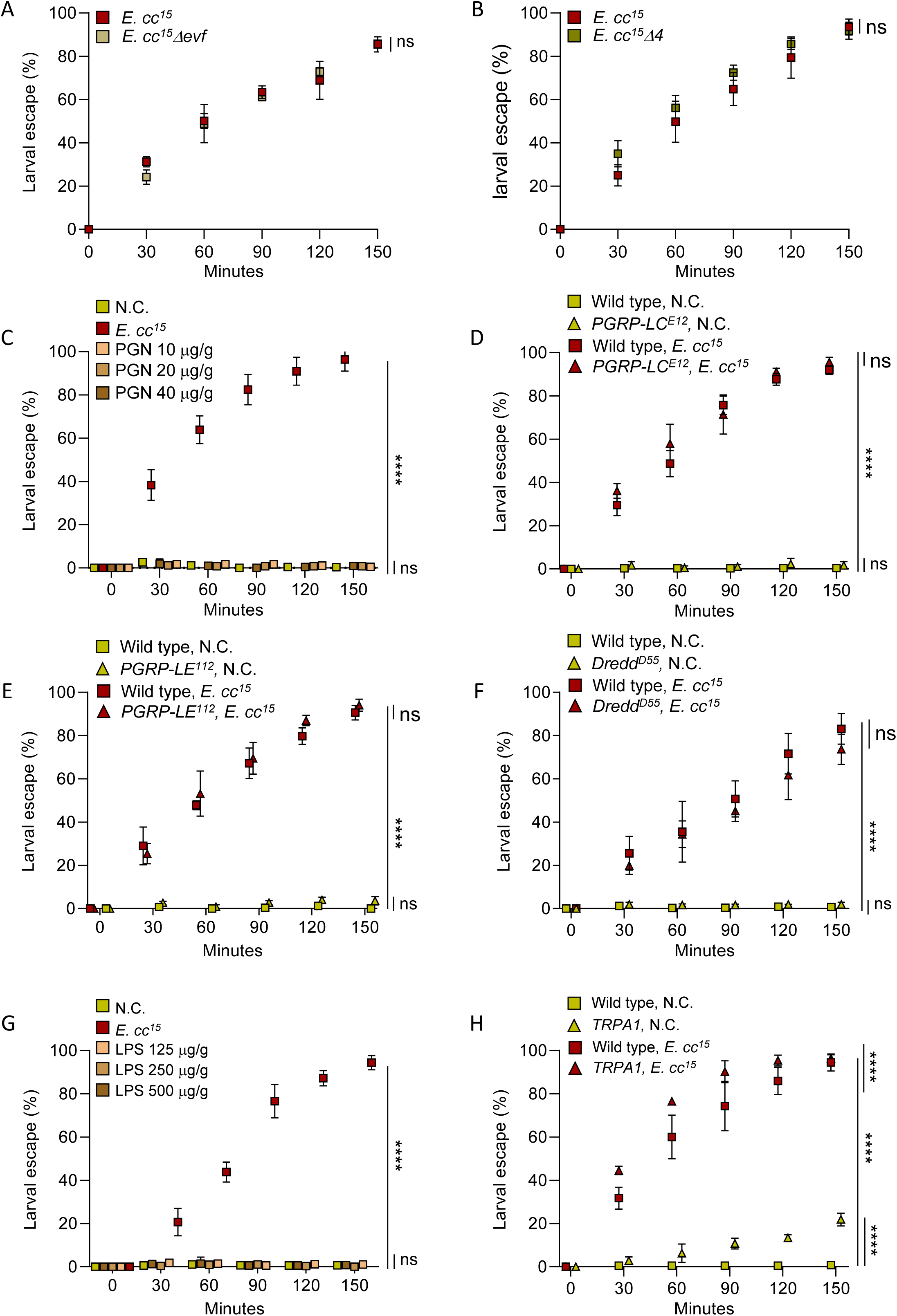
*E. cc^15^*-induced larval dispersal occurs independently of major bacterial factors, IMD immune signaling, and TRPA1 nociception. (A) Larval response to yeast contaminated with wild-type *E. cc^15^* or the Evf-deficient mutant (*E. cc^15^ Δevf*). **(B)** Escape behavior induced by wild-type or uracil-deficient *E. cc^15^* (*E. cc^15^ Δ4*). **(C)** Larval dispersal on food supplemented with increasing concentrations of purified PGN (10, 20, 40 μg/mL). **(D–F)** Dispersal of IMD pathway mutants: *PGRP-LC^E12^* (D), *PGRP-LE^112^*(E), and *Dredd^D55^* (F) on uncontaminated yeast and yeast mixed with *E cc^15^*. **(G)** Escape response to yeast supplemented with purified LPS (125, 250, 500 µg/mL). (F) *TRPA1* mutant larvae behavior on uncontaminated and *E. cc^15^*-contaminated food. Statistical analyses were performed using a mixed-effects model (REML) with time and condition as fixed factors, (ns = not significant; *p < 0.05; **p < 0.01; ***p < 0.001; ****p < 0.0001). Comparisons between selected conditions are shown.

The role of uracil, a microbe-derived metabolite known to activate DUOX-dependent epithelial defenses in adults [16], was then examined. Comparable levels of dispersal were observed when larvae were exposed to wild-type or uracil-deficient *E. cc^15^* (*E. cc^15^ Δ4*) [28], showing that uracil is also dispensable (Fig 3B).

The potential involvement of two conserved Gram-negative cell wall components was next evaluated. Although *E. cc^15^*-derived PGN is a strong activator of IMD signaling [13,15] and can induce aversive feeding responses in adult flies [29], supplementation of yeast with increasing concentrations of purified PGN did not induce dispersal (Fig 3C). In agreement, larvae lacking the PGN receptors PGRP-LC or PGRP-LE [30,31], or the downstream IMD effector Dredd [32], dispersed from *E. cc^15^*-contaminated food, demonstrating that PGN detection and IMD signaling are not required (Fig. 3D–F). While LPS is known to activate TRPA1-dependent avoidance pathways in adult flies [22], the addition of purified LPS to yeast failed to trigger larval dispersal (Fig 3G). Consistently, loss of TRPA1 function did not impair dispersal on *E. cc^15^*-contaminated yeast (Fig 3H).

Taken together, these findings show that larval dispersal from *E. cc^15^*-contaminated food is not mediated by individual bacterial factors such as Evf, uracil, PGN, or LPS, nor does it depend on canonical IMD-dependent detection or TRPA1-mediated sensing of noxious cues. Instead, the behavior is likely driven by the integration of *E. cc^15^* -derived signals through alternative sensory pathways.

### Aversive gustatory receptor Gr33a mediates larval dispersal from *E. cc^15^*-contaminated food

In *Drosophila*, non-volatile aversive cues are detected by gustatory receptor neurons (GRNs) expressing specific gustatory receptors (GRs) [33–35]. Among the larval neurons, Gr33a and Gr66a are co-expressed in bilateral pairs located primarily in the terminal organ (TO) and in pharyngeal sensilla, which together constitute the core aversion-sensitive GRN population [36]. To evaluate their functional requirement in *E. cc^15^*-induced dispersal, Gr66a- and Gr33a-positive neurons were silenced by expressing Kir2.1 (*Gr66a-GAL4>UAS-Kir2.1* and *Gr33a-GAL4>UAS-Kir2.1*), hyperpolarizing these GRNs and preventing synaptic transmission [37]. This markedly reduced dispersal from *E. cc^15^*-contaminated food compared to controls, without affecting behavior on uncontaminated yeast (Fig 4A, S7A Fig). Thus, these aversive GRNs are essential for the escape response triggered by bacterial contamination.

**Fig 4.**
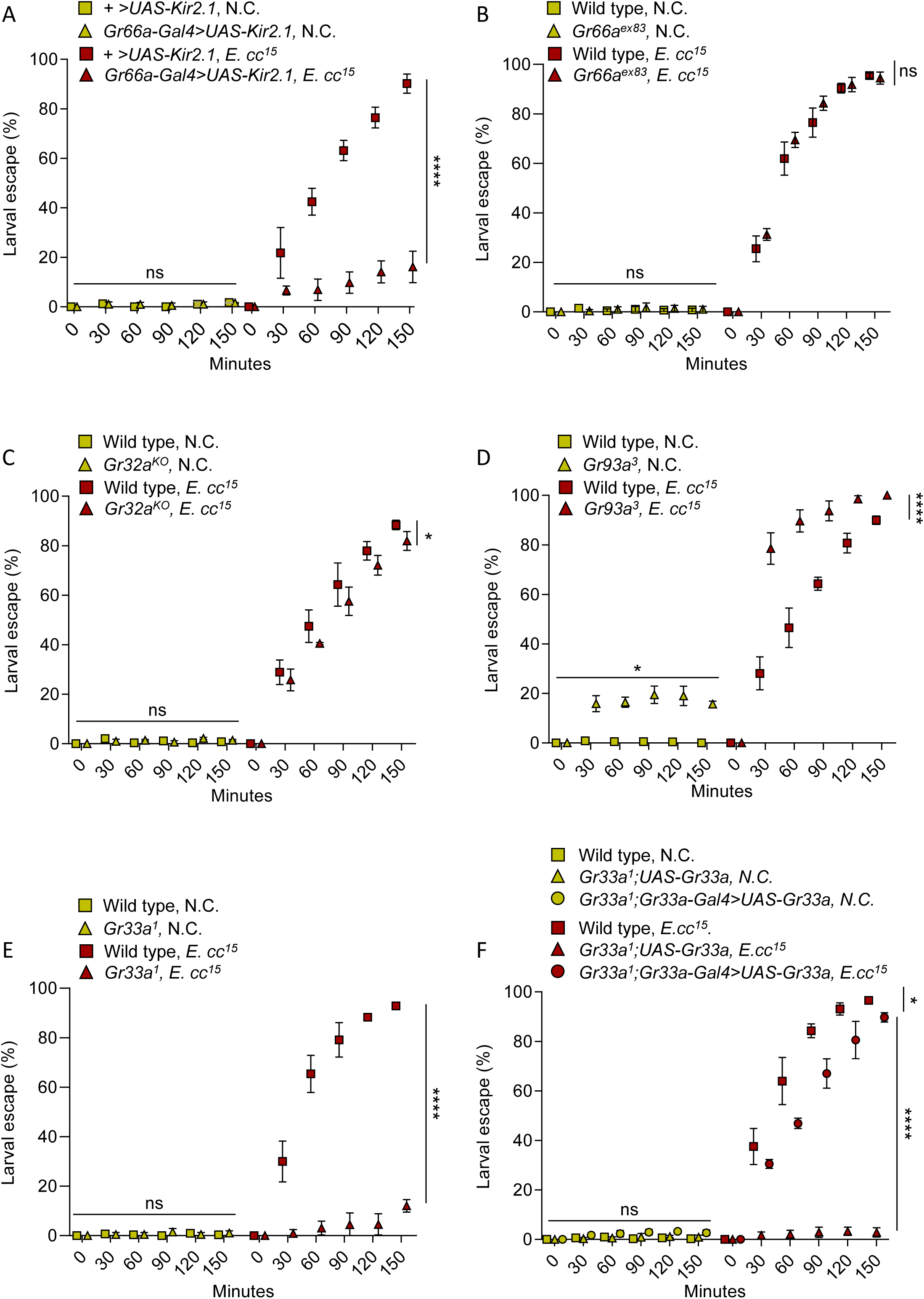
The gustatory receptor Gr33a mediates larval escape from *E. cc^15^-*contaminated food. **(A)** Larval dispersal following silencing of Gr66a-expressing neurons (*Gr66a-GAL4>UAS-Kir2.1*) on uncontaminated (N.C.) or *E. cc^15^*-contaminated food. **(B-E)** Escape response of wild type larvae and loss-of-function mutants *GR66a^ex83^* (B), *GR32a^KO^* (C), *GR93a^3^*(D) and *Gr33a^1^* exposed to uncontaminated and *E. cc^15^*-contaminated food. **(F)** Comparison of dispersal on control and *E. cc^15^*-containing food in wild-type, *Gr33a^1^; UAS-Gr33a,* and *Gr33a^1^; Gr33aGAL4>UAS-Gr33a* genotypes. Statistical analyses were performed using a mixed-effects model (REML) with time and condition as fixed factors, (ns = not significant; *p < 0.05; **p < 0.01; ***p < 0.001; ****p < 0.0001). Comparisons between selected conditions are shown. For all Figures, baseline behavioral characterization of parental GAL4 lines on *E. cc^15^*-contaminated food are provided in S Fig 14.

Within the Gr66a/Gr33a neurons, additional receptors linked to aversive contact behaviors, Gr32a and Gr93a, are expressed in partially overlapping subsets [38–40,36]. To determine their specific contributions, loss-of-function mutant alleles for *Gr33a* (*Gr33a¹*), *Gr66a* (*Gr66a^ex83^*), *Gr32a* (*Gr32a^KO^*), and *Gr93a* (*Gr93a³*) were analyzed. *Gr66a^ex83^* and *Gr32a^KO^* larvae dispersed at wild-type levels from contaminated food (Fig 4B–C), indicating that these receptors are not required. *Gr93a³* mutants showed heightened dispersal on contaminated food and mild nonspecific dispersal on clean yeast (Fig 4D), suggesting Gr93a modulates gustatory sensitivity rather than specifically mediating bacterial avoidance. By contrast, *Gr33a¹* larvae showed a strong dispersal defect on *E. cc^15^*-contaminated yeast, remaining on the substrate throughout the assay (Fig 4E). This phenotype was not attributable to defects in attraction or locomotion: when positioned at a distance from a food source, *Gr33a¹* larvae readily oriented toward and reached both clean and contaminated substrates, but, unlike controls, failed to withdraw after contacting *E. cc^15^*-contaminated yeast (S7B Fig). Reintroducing *Gr33a* expression (*Gr33a¹; Gr33a-GAL4>UAS-Gr33a*) restored dispersal to wild-type levels (Fig 4F). Together, these results identify Gr33a as required for dispersal from *E. cc^15^* contamination, highlighting its key role in contact-dependent detection of bacterial cues within the larval gustatory circuit.

### *orco–Or49a* olfactory and *Gr33a* gustatory circuits cooperate to mediate larval escape from *E. cc^15^*-contaminated food

Olfaction provides a critical sensory input that complements gustation in detecting aversive environmental cues [41]. To determine whether volatile compounds contribute to the dispersal behavior triggered by bacterial contamination, larvae carrying null mutations in the olfactory co-receptor *orco* (*orco¹* and *orco²*) were tested on *E. cc^15^*-contaminated yeast. While wild type and heterozygous *orco¹/+* larvae rapidly escaped from the contaminated substrate, anosmic *orco¹* mutants displayed a complete loss of this avoidance behavior (Fig 5A). Similar results were obtained with *orco²* mutants and *orco¹/orco²* transheterozygotes (S8A-B Fig). Consistently, blocking synaptic transmission in *orco*-expressing olfactory receptor neurons (ORNs) using the temperature-sensitive blocker *shibire^ts^* [37] (*orco-GAL4>UAS-shi^ts^*) markedly reduced dispersal from contaminated food (S8C–D Fig). These findings indicate that *orco*-dependent ORNs are essential for detecting volatile cues that trigger larval escape.

**Fig 5.**
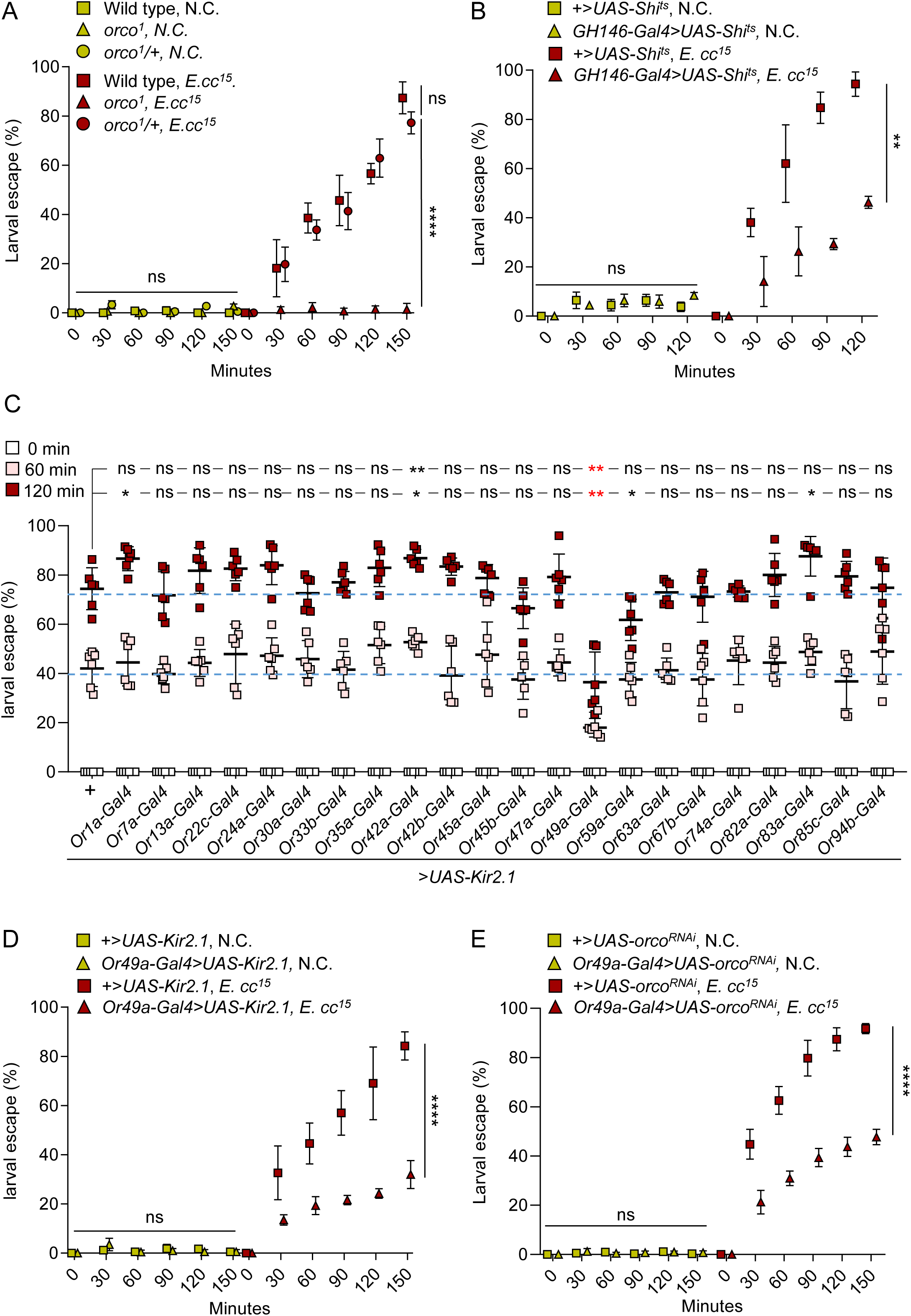
The orco–Or49a olfactory pathway mediates larval escape response to *E. cc^15^*-contaminated food. **(A)** Behavioral response of wild type, *orco¹* mutant and *orco¹*/+ heterozygotes larvae on uncontaminated (N.C) and *E. cc^15^*-supplemented yeast. **(B)** Response of *GH146-GAL4 > UAS-Shi^ts^* larvae to yeast– *E. cc^15^*contaminated food at *restrictive* temperature (31°C). **(C)** Escape induced by *E. cc^15^* at 60 and 120 min after silencing individual larval olfactory neuron classes (*OrX-GAL4 > UAS-Kir2.1*). Data are presented as means ± SD. Individual squares represent biological replicates (each consisting in groups of ≥ *40* larvae. ***(D)*** Larval dispersal *Or49a-GAL4>UAS-Kir2.1* larvae on uncontaminated or *E. cc^15^*-contaminated food. **(E)** Escape behavior of *Or49a-GAL4 > UAS-orco^RNAi^* larvae on *E. cc^15^*-contaminated food. Statistical analyses were performed using a mixed-effects model (REML) with time and condition as fixed factors for panels A, B, D, and E (ns = not significant; *p < 0.05; **p < 0.01; ***p < 0.001; ****p < 0.0001). Comparisons between selected conditions are shown. Panel C was analyzed using the Mann–Whitney test (*p < 0.05; **p < 0.01; ns = not significant).

To further delineate the olfactory circuitry, the role of projection neurons (PNs) transmitting sensory input from peripheral ORNs was examined. *orco*-positive ORNs project their axons to specific glomeruli within the larval antennal lobe, where they synapse onto GH146-GAL4-labeled PNs [42]. Silencing these neurons with *shibires^ts^* strongly reduced dispersal on contaminated yeast, without affecting behavior on uncontaminated yeast (Fig 5B), demonstrating that *orco*-dependent PNs are essential for olfactory signal transmission driving dispersal.

GH146-positive PNs send axons to two higher-order processing centers: the lateral horn (LH), which mediates innate olfactory responses, and the mushroom body (MB), involved in learning and multisensory integration [43,44]. Blocking synaptic transmission in Kenyon cells, the intrinsic MB neurons, using *OK107-GAL4 > UAS-shibire^ts^* [45] had no effect on larval escape from contaminated food (S9 Fig), suggesting that the MB is dispensable and that the rapid dispersal response relies on innate olfactory processing.

Next, the role of individual odorant receptors (Ors) was investigated. The larval olfactory system contains 21 ORNs in the dorsal organ (DO), each expressing a specific Or together with orco [46]. Selective suppression of Orco-dependent ORN subsets via Kir2.1 hyperpolarization (*OrX-GAL4 > UAS-Kir2.1*) showed that inhibition of *Or49a*-positive neurons specifically and reproducibly reduced dispersal, without affecting behavior on uncontaminated yeast (Fig 5C–D). To rule out locomotor or orientation deficits, *Or49a-GAL4 > UAS-Kir2.1* larvae were placed away from the food source; they oriented toward both uncontaminated and contaminated yeast but exhibited reduced escape frequency after contact with contaminated food (S10 Fig), consistent with impaired escape initiation.

*Or49a* is expressed in one neuron in the DO and another in the TO [46] (S11 Fig.). Targeted knockdown of *orco* in *Or49a*-expressing neurons (*Or49a-GAL4 > UAS-orco^RNAi^*) significantly decreased dispersal (Fig 5E), confirming that Or49a–Orco complexes in the DO mediate detection of volatile bacterial cues triggering escape. Double-driver expression analysis (*Gr33a-GAL4; Or49a-GAL4 > UAS-GFP*) showed non-overlapping *Or49a-* and *Gr33a*-expressing neurons in the TO, ruling out a shared circuit (S11 Fig).

Together, these findings reveal that larval escape from *E. cc^15^*-contaminated food depends on a cooperative multisensory network integrating gustatory and olfactory inputs: *Gr33a*-dependent neurons detect contact-dependent bacterial cues, while Or49a–Orco signaling mediates perception of volatile compounds, jointly driving rapid dispersal.

### Larval behavioral response to *E. cc^15^*-contaminated food modulates survival, development, and pathogen dissemination

The preceding results show that after an initial non-selective exploration of food, larval dispersal from *E. cc^15^*-contaminated substrates is triggered by the integration of gustatory and olfactory cues, limiting pathogen exposure and raising the question of how this behavior affects larval fate. To assess the consequences of this exposure-minimizing response, second-instar larvae were placed on either uncontaminated yeast or yeast contaminated with *E. cc^15^* or *E. coli*, and their growth and development were monitored.

After 24 h, larvae maintained on uncontaminated or *E. coli*–contaminated food exhibited comparable growth (Fig. 6A–B). Consistent with this, both groups progressed normally through development, with similar pupation and adult emergence rates (Fig 6C–E and S12 Fig). In contrast, larvae exposed to *E. cc^15^* were found outside the contaminated substrate, showed no significant growth (Fig 6A-B), and rarely pupated, with almost no adult emergence (Fig 6C-E and S12 Fig). These findings indicate that *E. cc^15^* exposure induces dispersal and persistent avoidance, which under restrictive conditions ultimately results in developmental failure and death.

**Fig 6.**
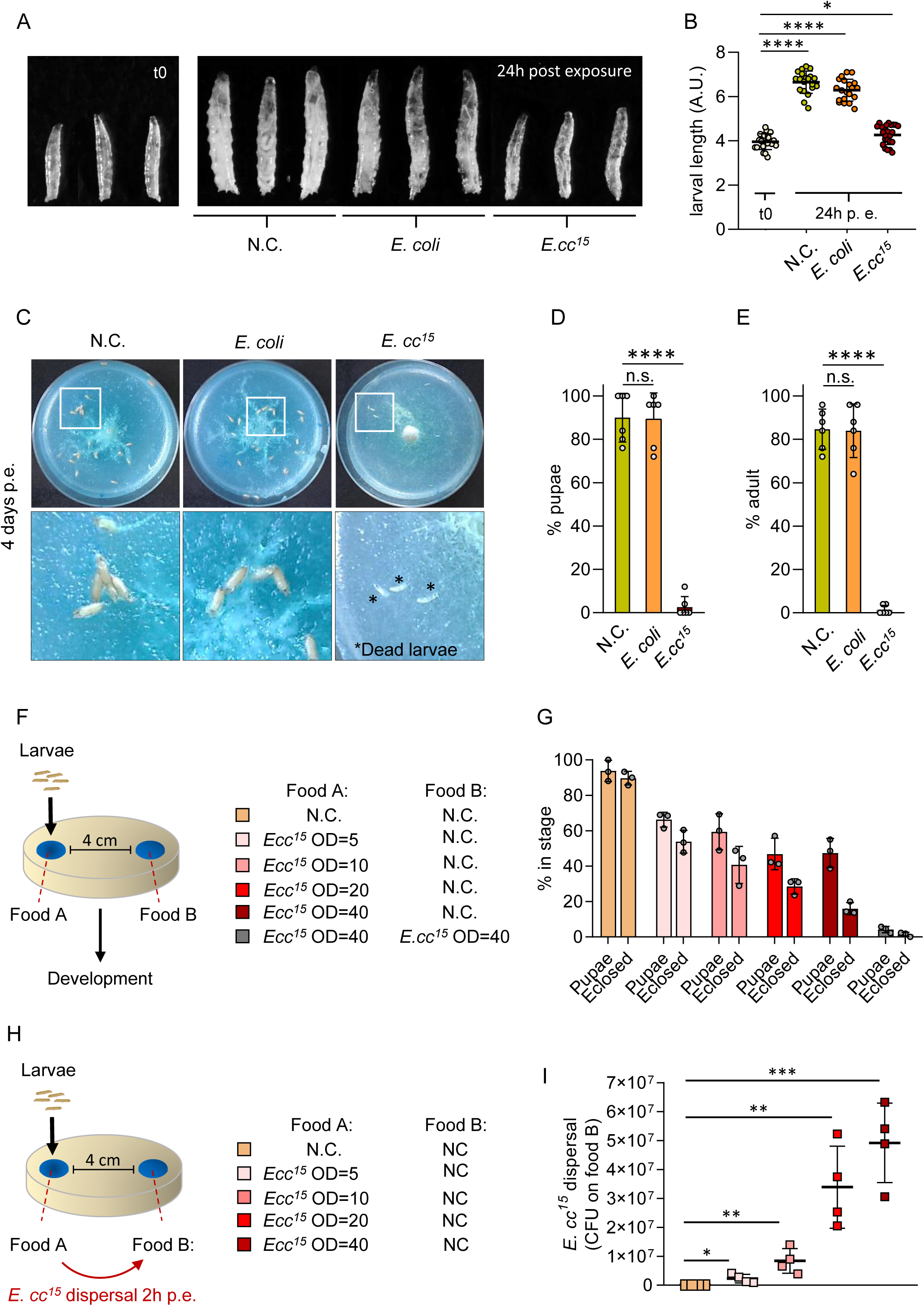
Exposure to *E. cc^15^*-contaminated food impairs larval development and promotes bacterial dissemination. **(A)** Representative images of larvae at 0 h and 24 h after exposure to control food (N.C.) or food supplemented with *E. coli* or *E. cc^15^*. **(B)** Quantification of larval body lengths at 0 h and 24 h following exposure to control or contaminated food (arbitrary units). Data are presented as means ± SD. Individual points represent individual larvae (n ≥ *20).* **(C)** Representative images illustrating pupal formation after exposure to control or contaminated food; boxed regions are shown at higher magnification below. **(D–E)** Proportion of larvae that reached the pupal stage (D) or successfully eclosed as adults (E) after exposure to control or contaminated food. Data are presented as means ± SD. Individual points represent biological replicates (each consisting of *25* larvae). **(F)** Schematic of the assay used to evaluate developmental outcomes of larvae that relocate to uncontaminated food after an initial exposure to *E. cc^15^*-contaminated substrate. **(G)** Proportion of larvae previously exposed to *E. cc^15^* that reached the pupal stage or eclosed after relocating to clean food (N.C). Data are presented as means ± SD. Individual points represent biological replicates (each consisting of *groups of* ≥ *35* larvae larvae). **(H)** Schematic of the bacterial dissemination assay. **(I)** *E. cc^15^* retrieved (CFU) from initially uncontaminated food (N.C) two hours after larvae were exposed to contaminated substrate. Data are presented as means ± SD. Individual squares represent biological replicates. Statistical analyses were performed using the Mann–Whitney test for panels B, D and E (*p < 0.05; **p < 0.01; ns = not significant). Panel G was analyzed using Chi-square tests that demonstrate that *E. cc^15^* concentration impacts on the pupation of larvae and on emergence of adult flies (****p < 0.0001). Panel I was analyzed using unpaired t test (ns = not significant; *p < 0.05; **p < 0.01; ***p < 0.001).

To evaluate whether dispersal provides an advantage when alternative resources are available, larvae were placed on yeast contaminated with increasing bacterial loads or on uncontaminated yeast, with each initial patch positioned at a distance from a second, clean food source (Fig 6F). After 24 hours, larvae initially placed on uncontaminated yeast largely remained on the original patch, indicating spatially restricted foraging in the absence of pathogen cues (S13A-B Fig). In contrast, larvae initially exposed to *E. cc^15^* consistently relocated to the uncontaminated patch at all bacterial concentrations, demonstrating a directed refuge-seeking behavior and minimal re-entry into contaminated food (S13B Fig). Developmental analysis revealed a dose-dependent impairment linked to the bacterial load, with pupation and eclosion rates significantly lower than those of larvae initialy placed on uncontaminated food (Fig 6G). These results suggest that access to an uncontaminated refuge partially mitigates the detrimental effects of initial *E. cc^15^* exposure.

Although this behavior provides only partial protection to the host under these experimental conditions, it may simultaneously facilitate pathogen dispersal. Similar host-mediated transmission has been described in adult *Drosophila* melanogaster, which can act as a natural vector of *E. cc^15^* between plant hosts [11,12]. To determine whether larval dispersal promotes bacterial spread, larvae were exposed to contaminated yeast at increasing bacterial loads, with each patch positioned at a distance from an uncontaminated food source (Fig 6F). Within two hours of exposure, larvae were consistently detected on the clean substrate, with the number of relocated individuals increasing with the initial bacterial load (S13C Fig). Viable *E. cc^15^* cells were recovered from these previously uncontaminated patches, with abundance correlating with both the number of migrating larvae and the initial bacterial load (Fig 6H).

Together, these findings indicate that the behavioral response elicited by *E. cc^15^*-contaminated food, including initial approach, contact, and directed withdrawal, not only limits host exposure but also enhances bacterial dissemination, suggesting that larval *Drosophila* can serve as an effective vector contributing to the environmental spread of this entomopathogen.

## DISCUSSION

Animals developing in microbe-rich environments often rely on early behavioral defenses to limit pathogen exposure before immune responses become engaged [19,47,48]. Here, we show that *Drosophila melanogaster* larvae display a rapid and robust escape behavior when encountering food contaminated with *E. cc^15^*. This response, which has not been previously described for this natural pathogen, expands the repertoire of known larval behavioral defenses and underscores the importance of ecological context in shaping pathogen detection.

Together, our findings identify a multisensory behavior that limits further exposure to contaminated substrate while concurrently promoting bacterial dissemination, revealing an unexpected ecological interaction between *Drosophila* larvae and *E. cc^15^*.

### Larval escape is triggered under specific ecological and bacterial metabolic conditions

Behavioral responses to pathogens have been far less explored in *Drosophila* larvae than in adults. Previous work demonstrated that larvae can abandon food contaminated with *Pseudomonas entomophila*, but this response is slow, state-dependent, and influenced by the physiological condition of the animal [49]. Under those experimental conditions, conducted on nutrient-poor apple substrate, larvae did not avoid *E. cc^15^* [49]. In agreement, we confirmed that stationary-phase *E. cc^15^* placed on agarose-based medium does not induce dispersal. Our results now demonstrate that larval escape requires that *E. cc^15^*be metabolically active, a condition achieved when stationary cells, which are physiologically quiescent and characterized by minimal metabolic activity, are provided access to nutrients that allow rapid regrowth. This requirement emphasizes a critical and previously overlooked point: the metabolic state of the bacteria strongly conditions their detectability and behavioral impact on the host. Dispersal becomes detectable shortly after regrowth begins, and the capacity of a given preparation to trigger avoidance is retained when the exposed patch is assayed days later, suggesting that the cues produced during regrowth persist on the substrate. Although the specific bacterial metabolites that trigger escape remain to be identified, soft-rot *Erwinia* species are well documented to secrete plant cell wall–degrading enzymes and other virulence-associated factors preferentially during metabolically active, quorum sensing–dependent growth phases, whereas these activities decline as cultures enter stationary phase [50–53]. These observations support the idea that the cues triggering larval escape are produced during lag and/or exponential growth phases, but not in stationary phase.

Importantly, under our experimental conditions, the escape response is elicited by *E. cc^15^* but not by non-pathogenic *Enterobacteriaceae* such as *E. coli*, consistent with the ability of larvae to develop on bacteria-rich substrates in natural environments. Whether other natural bacterial pathogens of *Drosophila*, such as *Serratia marcescens* or *Pseudomonas entomophila*, can similarly trigger rapid escape when metabolically active remains an open question; given the strong induction of virulence programs and volatile metabolites during growth of these species [8,54,55], such behaviors may be more widespread than currently appreciated.

### Larval escape emerges from the integration of gustatory and olfactory cues

Our findings reveal that larval escape from *E. cc^15^*relies on the cooperation of two distinct chemosensory modalities: gustation and olfaction. To our knowledge, this is the first demonstration of a rapid, multisensory escape behavior specifically triggered by a natural pathogen in *Drosophila* larvae. The gustatory receptor Gr33a, a broad aversion receptor associated with bitter and toxic compound detection [34,38–40], is essential for the response. Although Gr33a often functions together with Gr66a in adult taste circuits, our data indicate that Gr33a acts here independently of Gr66a, pointing to a Gr33a-dependent mechanism of danger detection. The requirement for contact-dependent gustatory input is consistent with the presence of at least one non-volatile metabolite produced during bacterial growth that signals substrate contamination.

In parallel, an olfactory component is required. The odorant receptor Or49a, previously implicated in the detection of parasitoid-derived iridomyrmecin [56], is also essential for dispersal. This finding indicates that Or49a is required for detecting a volatile cue produced by *E. cc^15^*, distinct from iridomyrmecin, highlighting the functional versatility of this olfactory channel. The involvement of Or49a aligns with growing evidence that the larval olfactory system, though anatomically simple, contains dedicated circuits for detecting ecologically salient threats [56,57].

Strikingly, disruption of either the gustatory or the olfactory pathway abolishes escape. This demonstrates that larvae rely on a cooperative multisensory process in which volatile and contact-dependent cues provide complementary information about substrate danger. Whether these cues correspond to different chemical forms of the same compound or represent distinct metabolites remains unknown. Nevertheless, the fact that two independent chemosensory modalities are each required supports the view that the larval response depends on the integration of multiple bacterial signals.

The neural mechanisms by which larvae combine gustatory and olfactory cues remain poorly characterized. By silencing GH146-positive projection neurons, which innervate the larval antennal lobe and relay olfactory information to higher brain centers, we show that the escape response requires transmission through the major olfactory projection pathways [43,57]. These projection neurons send outputs to both LH and the mushroom body MB [58–60], yet silencing Kenyon cells had no effect, suggesting that associative MB circuits are not required. Taste information from Gr33a-expressing neurons is relayed to the subesophageal zone (SEZ), the primary gustatory processing center in larvae [36,38,40]. While the output pathways of the SEZ have been only partially mapped, they clearly connect gustatory processing to multiple higher-order protocerebral regions. Although our data do not pinpoint where gustatory and olfactory streams converge, the combined requirement for GH146-positive projection neurons and Gr33a-expressing neurons, together with the rapid, stereotyped, and decision-like nature of the behavior, supports the view that escape is likely routed through non-associative circuits consistent with innate valence pathways. This interpretation aligns with the established role of the LH in mediating hardwired responses to ecologically salient stimuli, whereas the mushroom body MB is generally required for learned aversions [61,62].

Importantly, this behavioral paradigm provides a tractable model for dissecting how taste and odor signals are integrated to guide ecologically relevant decisions. The genetic accessibility of the larva, together with the ability to manipulate both sensory modalities independently, makes this system a powerful framework for mapping the cellular and circuit mechanisms through which bacterial danger cues are transformed into behavioral action.

### A protective behavior that simultaneously promotes pathogen dissemination

Larval escape provides an opportunity to reduce exposure to pathogenic bacteria by enabling migration toward uncontaminated food. Developmental analyses show that access to a refuge partially mitigates the deleterious effects of initial exposure to *E. cc^15^*, with higher pupation and eclosion rates compared to larvae that fail to locate an alternative food source. Thus, the behavior can confer a protective benefit, but only under ecological conditions where clean patches are accessible. These findings underscore that the adaptive value of escape is conditional rather than absolute, depending on the spatial structure and heterogeneity of the environment.

Escape, however, has a second ecological consequence: it promotes *E. cc^15^* dispersal. Viable bacteria were consistently recovered from previously uncontaminated patches reached by dispersing larvae, indicating that larval movement can mechanically spread bacterial cells. Comparable host-mediated transmission processes are well documented for plant-associated pathogens, in which insect mobility frequently contributes to pathogen dissemination [63]. Notably, adult *Drosophila* are natural vectors of *Erwinia* species [11,12], suggesting that both larval and adult stages may contribute to the environmental spread of this bacterium.

These observations raise the possibility that larval escape could simultaneously benefit both host and pathogen: larvae reduce exposure by abandoning contaminated food, while *E. cc^15^* is mechanically transported to new substrates through their movement. Whether this dual outcome reflects an evolved manipulation strategy or simply the incidental exploitation of a pre-existing host defense remains unclear. Our data do not support active behavioral manipulation by the pathogen; however, the fact that escape is triggered only during active bacterial growth suggests that metabolic pathways linked to virulence or microbial competition may inadvertently promote larval dispersal. More broadly, the coexistence of host-protective and pathogen-promoting consequences illustrates how interactions between microbial cues and host decision-making can generate ecological feedbacks that influence pathogen spread.

### Conclusions and perspectives

This work establishes *E. cc^15^*-driven escape as a powerful model to investigate larval behavioral immunity, multisensory integration, and host–microbe ecological interactions. Identifying the metabolites produced by *E. cc^15^* that trigger dispersal will be an important next step and may uncover new classes of microbe-derived behavioral modulators. Broadening this approach to additional natural *Drosophila* pathogens will help determine how widespread rapid escape is as a defensive strategy across microbial threats.

More broadly, the conserved organization of chemosensory systems across dipterans suggests that the principles uncovered here may extend beyond *D. melanogaster*. Insights into how insects detect and avoid pathogen-associated cues may ultimately contribute to the development of environmentally friendly repellents or attractants based on microbial semiochemicals, offering potential avenues for the management of fruit- and fermentation-associated dipteran pests, which rely on similar ecological niches and chemosensory strategies.

## MATERIALS and METHODS

### Fly stocks and husbandry

The following *Drosophila melanogaster* strains were used in this study. The w1118 strain (no. 5905, Bloomington *Drosophila* Stock Center, BDSC) served as the reference wild-type strain in all main experiments unless otherwise indicated. Additional lines obtained from the BDSC included: *Canton-S* (no. 64349), *Oregon-R* (no. 25211), *PGRP-LE^112^* (no. 33055), *orco^1^* (no. 23129), *orco^2^* (no. 23130), *GH146-GAL4* (no. 91812), *OK107-GAL4* (no. 854), *UAS-Shibire^ts^*(no. 44222), *UAS-Kir2.1* (no. 6595), *Gr66a-GAL4* (no. 28801), *Gr33a-GAL4* (no. 57624), *Orco-GAL4* (no. 23292), *UAS-mCD4-Tomato* (no. 35841), *Gr66a^ex83^* (no. 25027), *Gr33a^1^* (no. 31427), *Gr93a^3^* (no. 27592), *Gr33a^1^*; *Gr33a-GAL4* (no. 31425), *Gr33a^1^; UAS-Gr33a-GAL4* (no. 31424), *Or1a-GAL4* (no. 9949), *Or7a-GAL4* (no. 23907), *Or13a-GAL4* (no. 9945), *Or22c-GAL4* (no. 9953), *Or24a-GAL4* (no. 9957), *Or30a-GAL4* (no. 9960), *Or33b-GAL4* (no. 9964), *Or35a-GAL4* (no. 9968), *Or42a-GAL4* (no. 9972), *Or42b-GAL4* (no. 9976), *Or47a-GAL4* (no. 9982), *Or49a-GAL4* (no. 9985), *Or59a-GAL4* (no. 9989), *Or63a-GAL4* (no. 9991), *Or67b-GAL4* (no. 9996), *Or74a-GAL4* (no. 23124), *Or82a-GAL4* (no. 23125), *Or83a-GAL4* (no. 23127), *Or85c-GAL4* (no. 23914), *Or94b-GAL4* (no. 23145), *TRPA1* (no. 26504) and *UAS-6xGFP* (no. 52262). The *yellow white* reference strain was kindly provided by Dr. B. Charroux (Aix-Marseille University, France). The *Dredd^D55^* and *Gr32a^KO^* mutant strains were generous gifts from Dr. B. Lemaire (EPFL, Switzerland) and Dr. J.R. Carlson (Yale University, USA), respectively. The *UAS-Orco^RNAi^* line (KK100825) was obtained from the Vienna *Drosophila* Resource Center (VDRC). The *PGRP-LC^E12^*mutant was described previously [64].

Flies were maintained at 25 °C on standard yeast/cornmeal medium under a 12 h light/12 h dark cycle. For 1 L of food, 8.2 g agar (VWR), 80 g cornmeal (Westhove Farigel maize H1), and 80 g yeast extract (VWR) were boiled for 10 min, cooled, and supplemented with 5.2 g methylparaben sodium salt (MERCK) and 4 mL 99% propionic acid (CARLO ERBA).

### Cultivation and heat inactivation of bacteria and yeast

The bacterial strains used included rifampicin-resistant *Erwinia carotovora carotovora 15* (*E. cc^15^*) [6], *E. cc^15^ Δevf* [27], *E. cc^15^ Δ4* [28], and *Escherichia coli* K-12 M4100, kindly provided by Dr. I. Gomperts Boneca (Institut Pasteur, France).

Single colonies were inoculated in 250 mL LB broth and grown overnight at 250 rpm (*E. cc^15^*at 30 °C; *E. coli* at 37 °C). Growth was monitored by OD_c_. Cultures were centrifuged at 3800 rpm for 15 min, and pellets were resuspended to the desired OD_600_ before mixing with baker’s yeast.

Heat-killed *E. cc^15^* was obtained by incubating pellets at 95 °C for 10 min followed by rapid cooling at −20 °C. Baker’s yeast (*S. cerevisiae*) was inactivated at 80 °C for 10–15 min and cooled to room temperature. Heat-killed bacteria and yeast were processed identically to live cells in subsequent assays.

### Behavioral assays

A 100 µL bacterial suspension was mixed with 0.5 g commercial baker’s yeast (∼0.4 mL). Final OD_600_ was calculated as: Final OD_600_=100 μL × Initial OD_600_ / 500 μL. A 100 µL drop of yeast–bacteria paste was placed on LB agar plates and incubated for 30 min, unless otherwise specified. Second-instar larvae (48 h at 25°C after egg laying) were washed in water and tested in groups of n ≥ 40 per replicate, unless otherwise stated. Each experiment included at least three technical replicates and was independently repeated at least three times.

In dispersal assays, larvae were placed directly on food; in avoidance and binary choice assays, larvae were positioned 4 cm from the food source(s). At each time point, larvae outside the food were counted under a ZEISS Stemi 508 stereomicroscope. At the end of each assay, the food was suspended in water and larvae counted after liquid removal. Escape percentages were calculated relative to the initial number of larvae.

Behavioral assays were performed at room temperature, unless otherwise specified, in darkness; illumination was applied only briefly during counting to minimize phototactic bias.

### Imaging of larval behavior

For time-point imaging, groups of >100 larvae were placed directly on the food source. Experiments were conducted in darkness, and white light was applied only briefly for image capture. Images were acquired using a Logitech HD Pro Webcam C920.

For movie generation, time-lapse imaging was performed using the same setup, controlled by Yawcam v0.8.0, capturing one frame every 2 s over 3 h 30 min. During acquisition, white-light intensity and direction were minimized to reduce phototactic responses. Image sequences were compiled into movies using Fiji (ImageJ v1.54p) and rendered at 24 fps. All images, including those used for time-point photography and movie generation, were processed with Photoshop CS6.

### Larval growth, development, and survival

Second-instar larvae were placed on control yeast or yeast mixed with *E. cc^15^*or *E. coli* (OD_600_ = 40). After 24 h, larvae were washed in phosphate-buffered saline (PBS), fixed in 70% ethanol, mounted, and imaged using a Zeiss Discovery Lumar V12 microscope. The size of ≥20 larvae per condition was quantified in arbitrary units using Fiji (ImageJ v1.54g) and compared to unexposed second instar larvae.

For development and survival assays, groups of 25 larvae per condition were monitored for pupation (day 4) and adult eclosion (day 7) following exposure to the food source. Counting was performed using a ZEISS Stemi 508 stereomicroscope, and images were acquired with a Logitech C920 camera.

### Assessment of bacterial dissemination by *Drosophila* larvae

To evaluate bacterial transfer from contaminated to uncontaminated food sources, groups of 25 larvae were placed on yeast mixed with *E. cc^15^* (OD_600_ = 5–40), located 4 cm from uncontaminated yeast. All conditions were performed in four technical replicates. After 2 h, the proportion of larvae that reached the uncontaminated yeast was quantified. The uncontaminated food was collected and homogenized in 2 mL PBS. A 1:1000 dilution was plated (200 µL) onto LB agar supplemented with rifampicin, incubated overnight at 30 °C, and colony-forming units (CFUs) counted to quantify viable *E. cc^15^* transferred by larvae.

### Bacterial components and culture media

Ultrapure peptidoglycan from *E. coli* (PGN-ECndi; InvivoGen, USA, cat. Tlrl-kipgn) and lipopolysaccharides (LPS) from *E. coli* O111: B4 (Sigma-Aldrich, cat. L2630) were used as bacterial components in experimental assays.

For the preparation of solid media, Luria–Bertani (LB) broth (Lennox formulation, Invitrogen, cat. L22897), agar (BD Bacto™, Becton Dickinson, cat. 214010), and high-purity agarose (NEEO Ultra, Euromedex, cat. 2267.4) were used. For liquid bacterial cultures, LB broth (Lennox formulation, Sigma-Aldrich, cat. L3022) was used. All reagents were handled according to the manufacturers’ instructions.

### Confocal imaging

Second instar larvae of the specified genotypes were heat-killed (60 °C, 10 s), mounted in Vectashield (Vector Labs, Cat. No. H-1200), and imaged using a Zeiss LSM 780 confocal microscope with a 20× air objective. Images were processed using Adobe Photoshop CS6.

### Data representation and statistical analyses

All graphical representations and statistical analyses were performed using GraphPad Prism 8. Larval escape dynamics were analyzed using a mixed-effects model (REML) with Time and Condition as fixed factors, including their interaction. The T0 time point was excluded because all conditions are identical at this stage. Post-hoc multiple comparisons were carried out using Tukey’s test. For all experiments, full raw datasets and complete statistical outputs are provided in S1 and S2 Data.

## AUTHOR CONTRIBUTIONS

Olivier Zugasti: Conceptualization, Methodology, Investigation, Formal analysis, Visualization, Writing – Original Draft Preparation and Data curation. Julien Royet: Conceptualization, Formal analysis, Writing – Original Draft Preparation and Funding acquisition.

## ACKNOWLEDGMENTS

We thank C. Léopold Kurz (Marseille Institute of Development Biology IBDM) for valuable scientific discussions and for critically reading the manuscript. We extend our appreciation to the Marseille Institute of Development Biology (IBDM) microscopy platform. This work was supported by CNRS, ANR Pepneuron (ANR-21-CE16-0027).

## SUPPLEMENTARY FIGURE LEGENDS

**S1 Fig.**
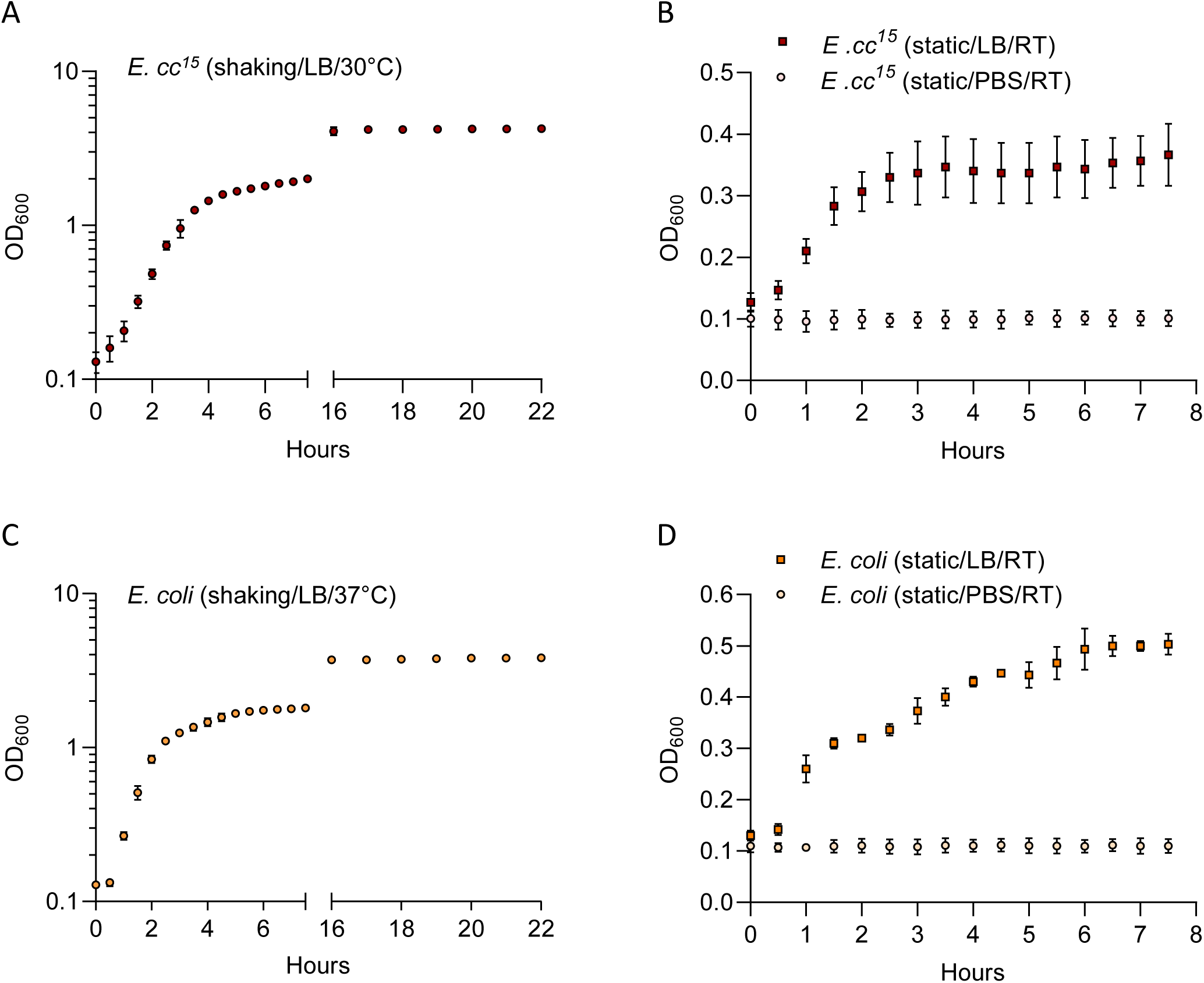
Growth dynamics of *E. cc^15^* and *E. coli* under distinct culture conditions. **(A)** Growth curve of *E. cc^15^* cultured in nutrient-rich LB under shaking conditions at 30 °C. **(B)** Growth or survival dynamics of stationary-phase *E. cc^15^* resuspended in fresh LB or PBS and maintained in static culture at room temperature. **(C)** Growth curve of *E. coli* cultured in nutrient-rich LB under shaking conditions at 37 °C. **(D)** Growth or survival dynamics of stationary-phase *E. coli* resuspended in fresh LB or PBS and maintained in static culture at room temperature. Data represent mean ± SD of three independent experimental replicates. For all supplementary figures, raw data and complete statistical analyses are provided in S2 Data.

**S2 Fig.**
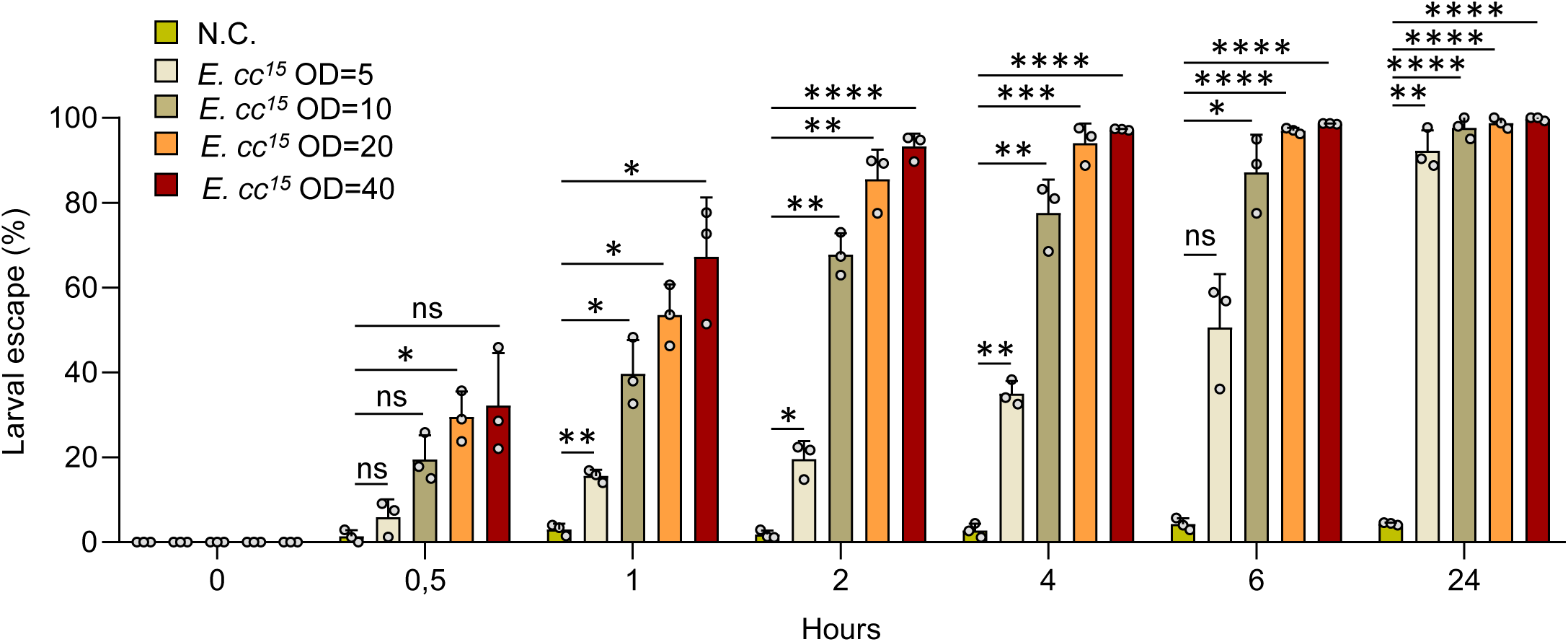
*E. cc^15^*triggers a sustained and dose-dependent escape behavior in Drosophila larvae. Quantification of second-instar larvae (w¹¹¹⁸) escape behavior from uncontaminated yeast (N.C.) or yeast contaminated with increasing concentrations of *E. cc^15^* (OD_600_ = 5, 10, 20, and 40) at multiple time points. For this and all subsequent supplementary figures, dispersal assay results are presented as mean ± SD of technical triplicates from a representative biological experiment. Each experiment was independently repeated at least three times with groups of ≥40 larvae per replicate, unless otherwise indicated. Statistical analyses were performed using two-way repeated measures ANOVA, followed by Tukey’s multiple comparisons test (**ns = not significant; *p < 0.05; **p < 0.01; ***p < 0.001; ****p < 0.0001). Comparisons between selected conditions are shown.

**S3 Fig.**
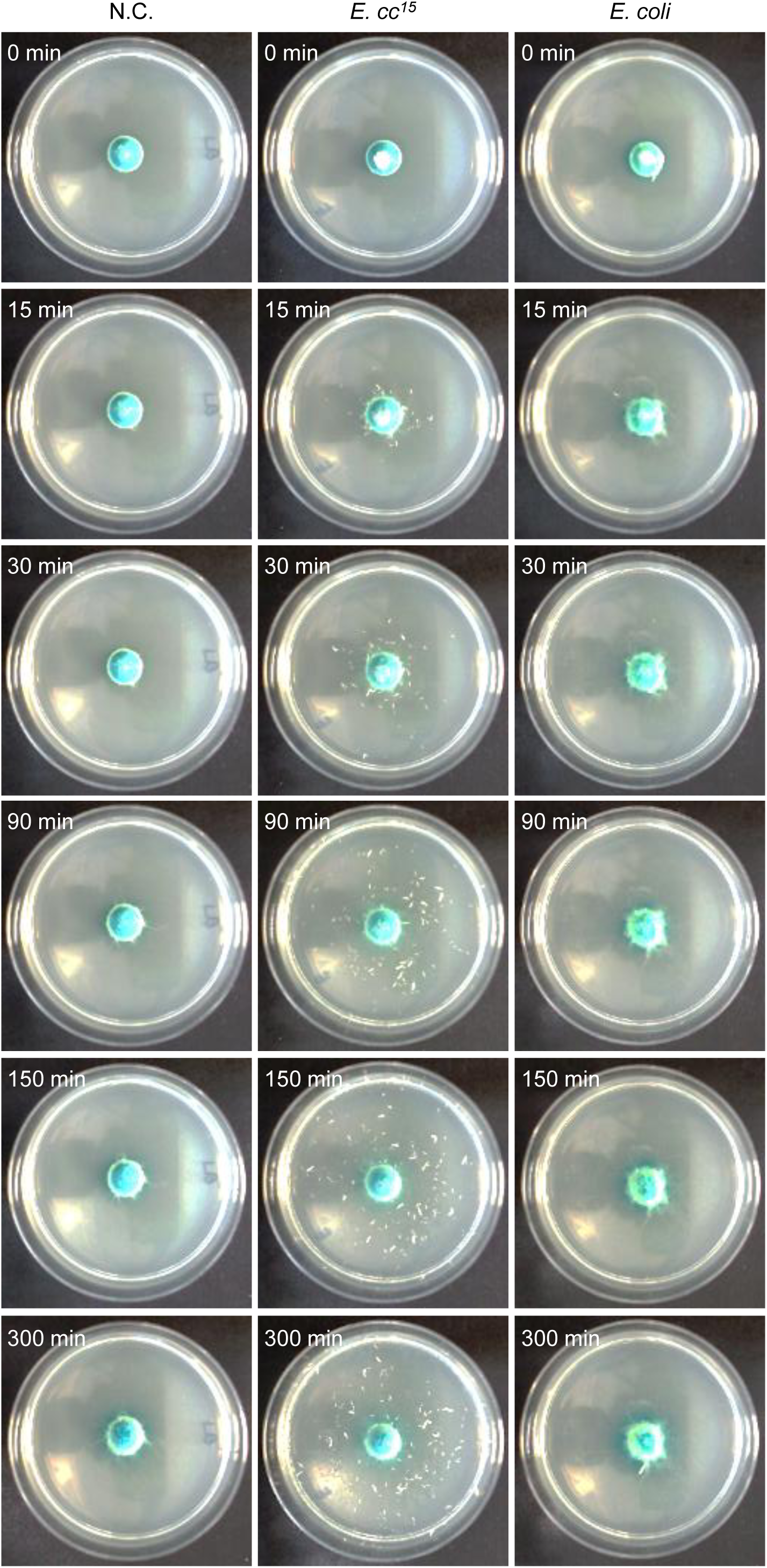
Time-course images of wild-type larvae on uncontaminated or bacterially contaminated yeast. Representative images of second-instar larvae (w¹¹¹⁸) at multiple time points, showing their positioning on either uncontaminated or *E. cc^15^***-** or *E. coli*-contaminated yeast (OD_600_ = 40). Yeast patches were deposited on nutrient-rich LB agar plates and **i**ncubated for 30 minutes at room temperature before imaging. In each condition, > 100 larvae were placed on the food source at t = 0. Images were captured using a standard webcam.

**S4 Fig.**
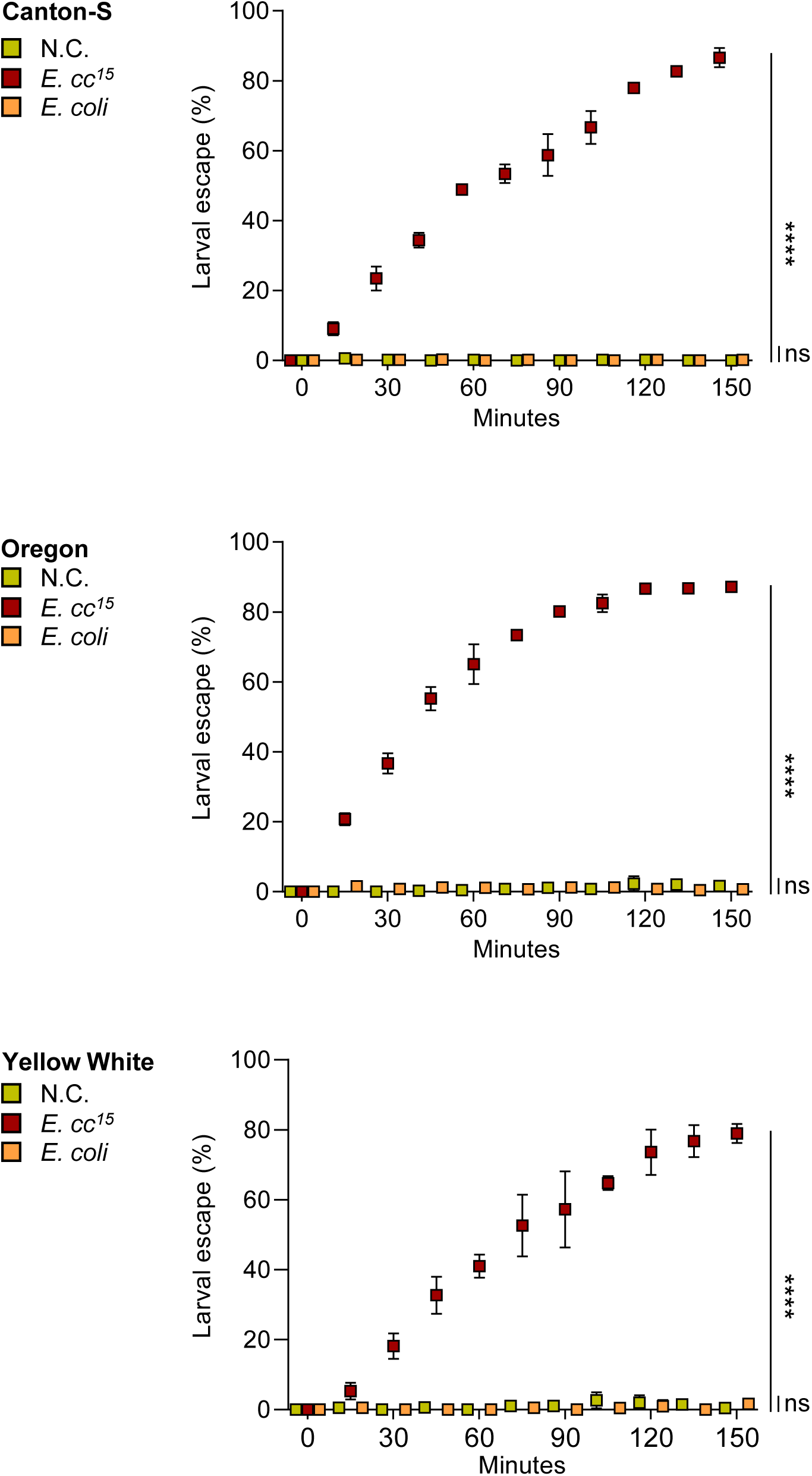
*E. cc^15^*-induced larval escape behavior is conserved across Drosophila melanogaster genetic backgrounds. Quantification of dispersal of second-instar larvae from Canton-S, Oregon-R, and Yellow white strains on either uncontaminated yeast (N.C.) or yeast containing *E. cc^15^* or *E. coli* at identical bacterial densities (OD_600_ = 40). Statistical analyses were performed using a mixed-effects model (REML) with time and condition as fixed factors (ns = not significant; *p < 0.05; **p < 0.01; ***p < 0.001; ****p < 0.0001). Pairwise comparisons between selected conditions are shown.

**S5 Fig.**
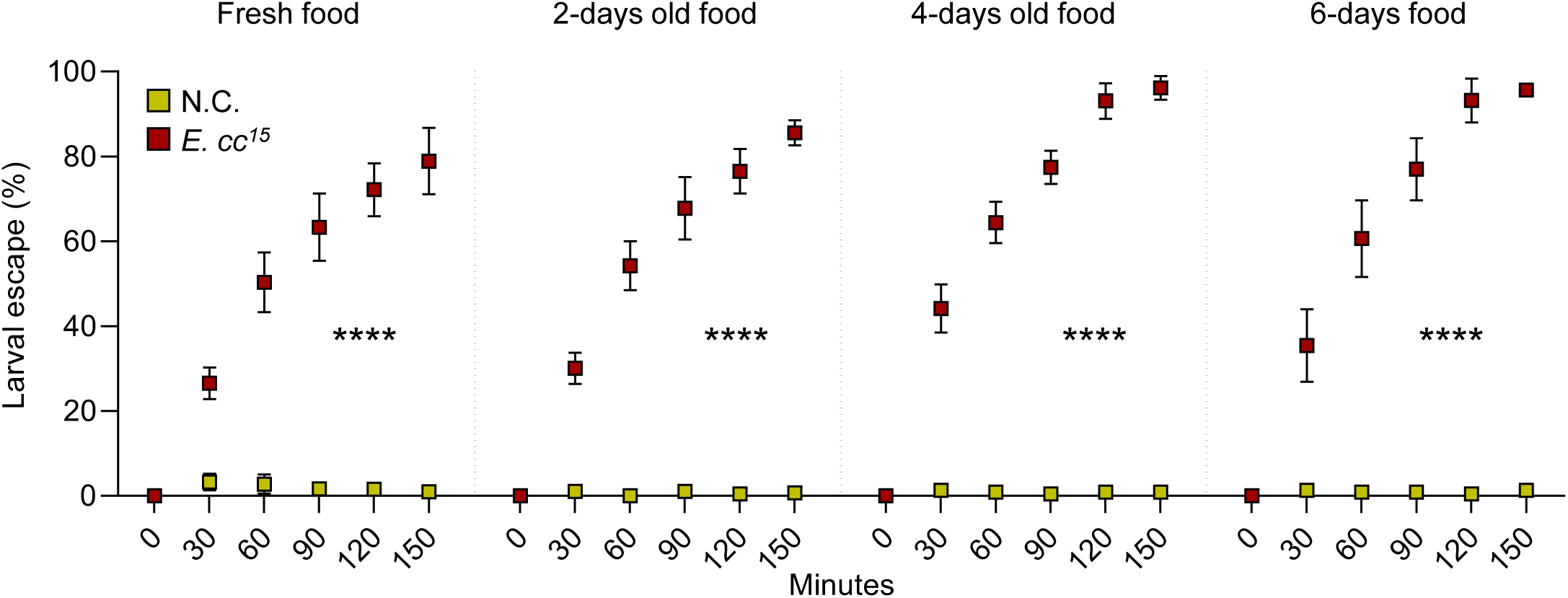
*E. cc^15^*-induced larval dispersal behavior is long-lasting and robust. Time-course quantification of second-instar *Drosophila* larvae (w¹¹¹⁸) escape behavior on freshly prepared *E. cc^15^*-contaminated yeast (OD₆₀₀ = 40) and on yeast–*E. cc^15^* mixtures maintained on LB agar plates for 2, 4, or 6 days. Statistical analyses were performed using a mixed-effects model (REML) with time and condition as fixed factors (ns = not significant; *p < 0.05; **p < 0.01; ***p < 0.001; ****p < 0.0001). Pairwise comparisons between selected conditions are shown.

**S6 Fig.**
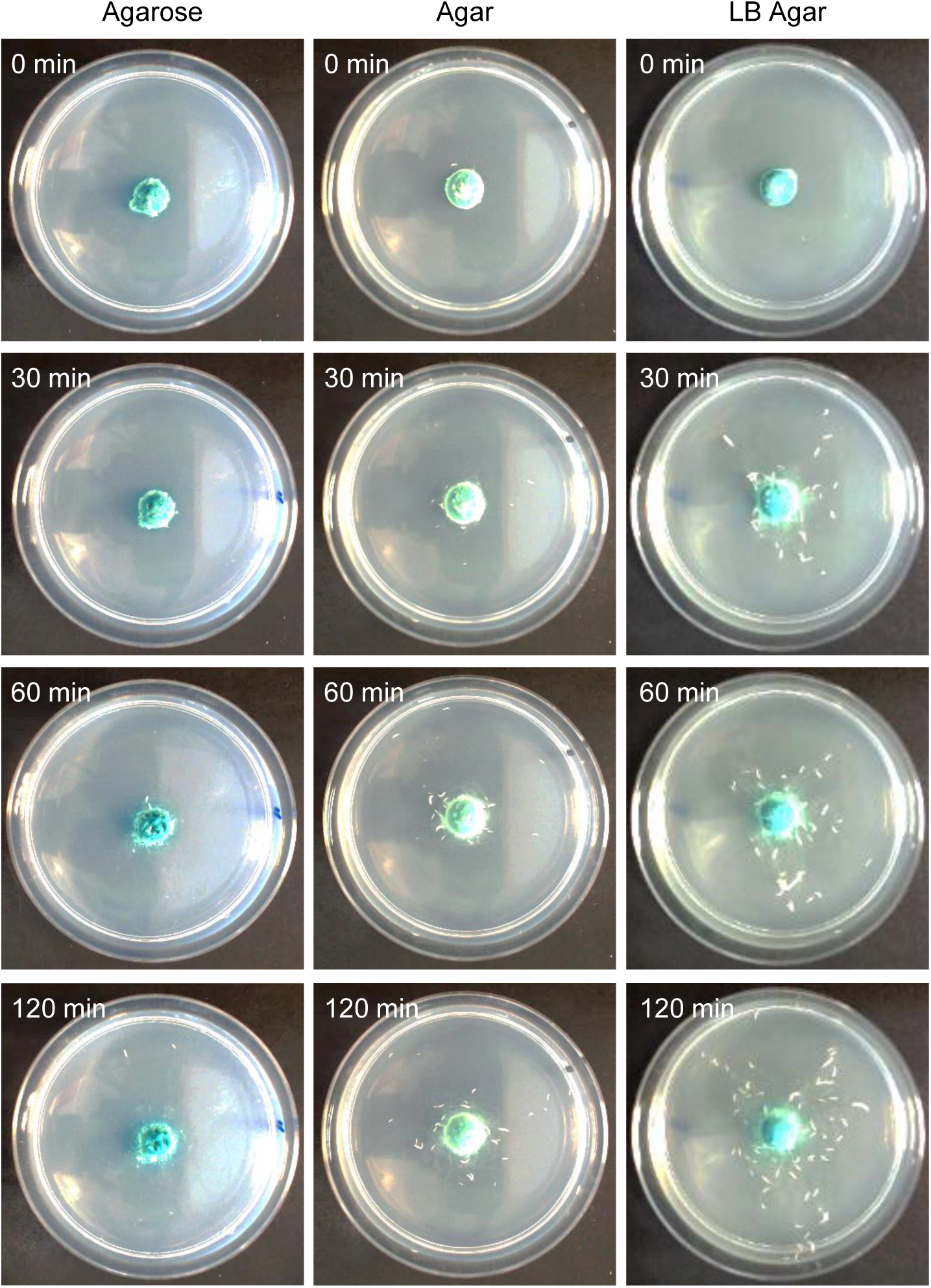
Nutrient availability in the substrate modulates *E. cc^15^*-induced larval dispersal. Representative time-course images of second-instar larvae (w¹¹¹⁸) on yeast–*E. cc^15^* mixtures (OD₆₀₀ = 40) placed on substrates with different nutrient content: nutrient-rich LB agar, nutrient-poor agar, and nutrient-free agarose. In each condition, >100 larvae were introduced onto the food at t = 0. Images were captured using a standard webcam.

**S7 Fig.**
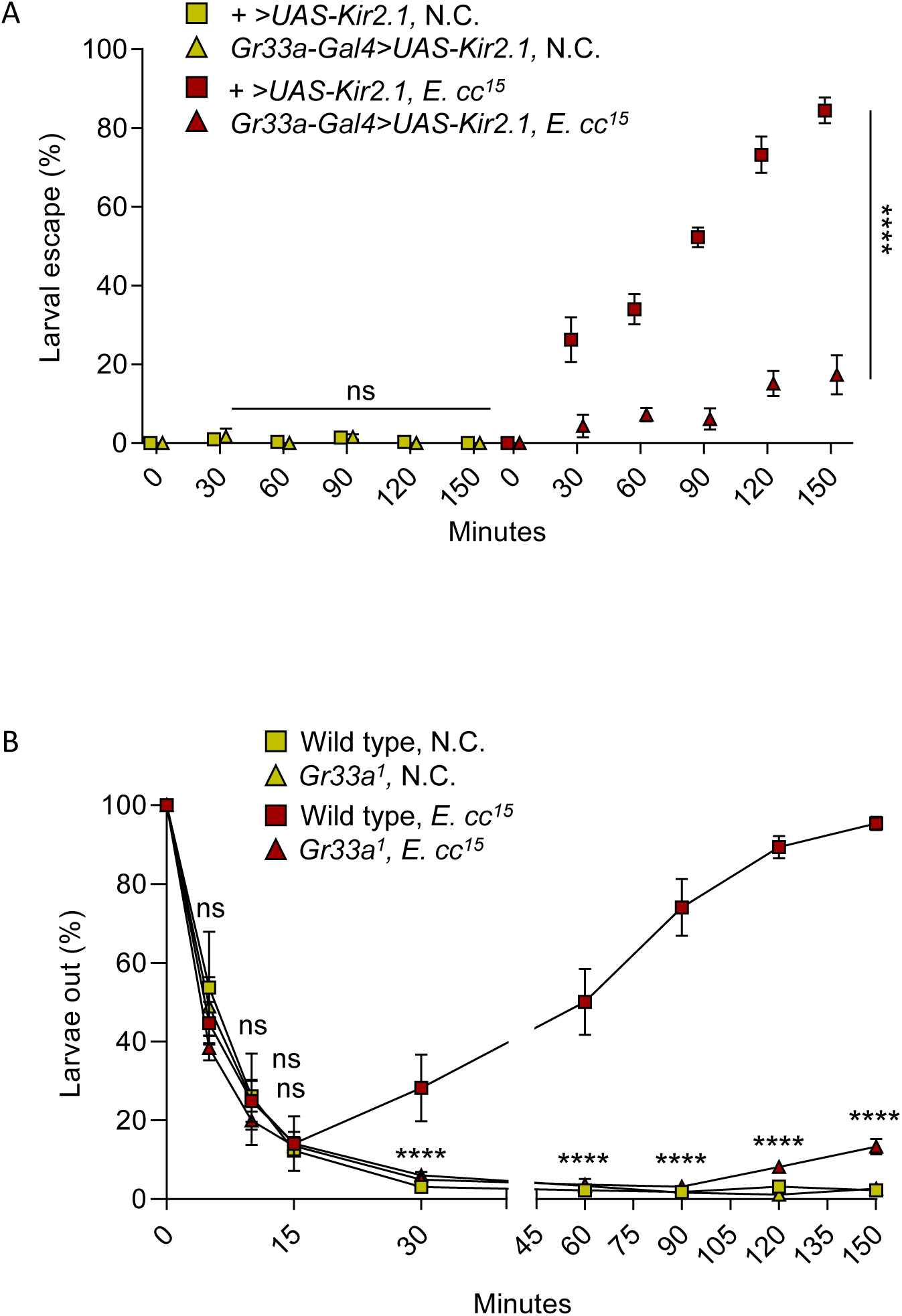
Silencing the *Gr33a* neuronal circuit suppresses *E. cc^15^*-induced escape without altering locomotion or attraction to nutritive cues. **(A)** Dispersal in control larvae *(+>UAS-Kir2.1*) and in larvae with silenced Gr33a-expressing neurons (*Gr33a-GAL4>UAS-Kir2.1*) on uncontaminated yeast (N.C.) or *E. cc^15^*-contaminated yeast. **(B)** Time-course quantification of wild-type and *Gr33a¹* mutant larvae located outside the food source when placed at a fixed distance from uncontaminated or *E. cc^15^*-contaminated yeast. Statistical analyses were performed using a mixed-effects model (REML) with time and condition as fixed factors; followed by Tukey’s post-hoc for-panel B comparing wild-type and *Gr33a¹* mutant larvae on *E. cc^15^*-contaminated food (ns = not significant; *p < 0.05; **p < 0.01; ***p < 0.001; ****p < 0.0001).

**S8 Fig.**
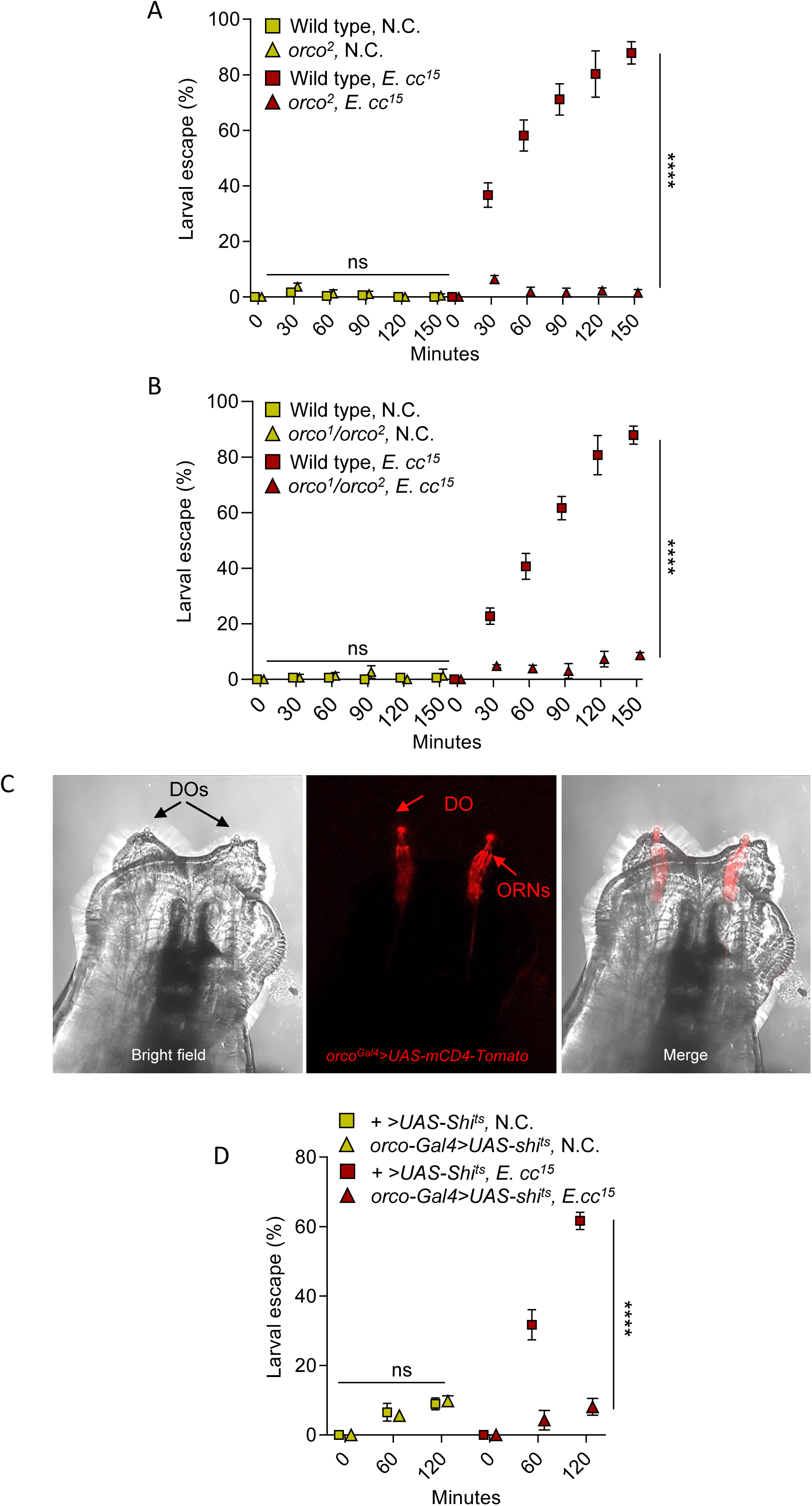
The odorant co-receptor orco is required for larval escape from *E. cc^15^*-contaminated food. **(A–B)** Time-course quantification of the escape behavior of wild-type, *orco²* mutant (A), and *orco¹/orco²* transheterozygous larvae (B) on uncontaminated yeast (N.C.) or *E. cc^15^*-contaminated yeast. **(C)** Confocal images showing, from left to right: bright-field view of dorsal organs (DOs, arrows); *Orco-GAL4>UAS-mCD4-Tomato* expression in olfactory receptor neurons (ORNs, arrows); merged image. **(D)** Escape behavior of control *(+>UAS-Shi^ts^*) and *Orco-GAL4>UAS-Shi^ts^* larvae exposed to **either** uncontaminated or *E. cc^15^*-contaminated yeast at the restrictive temperature (31 °C). Statistical analyses were performed using a mixed-effects model (REML) with time and condition as fixed factors (ns = not significant; *p < 0.05; **p < 0.01; ***p < 0.001; ****p < 0.0001). Pairwise comparisons between selected conditions are shown.

**S9 Fig.**
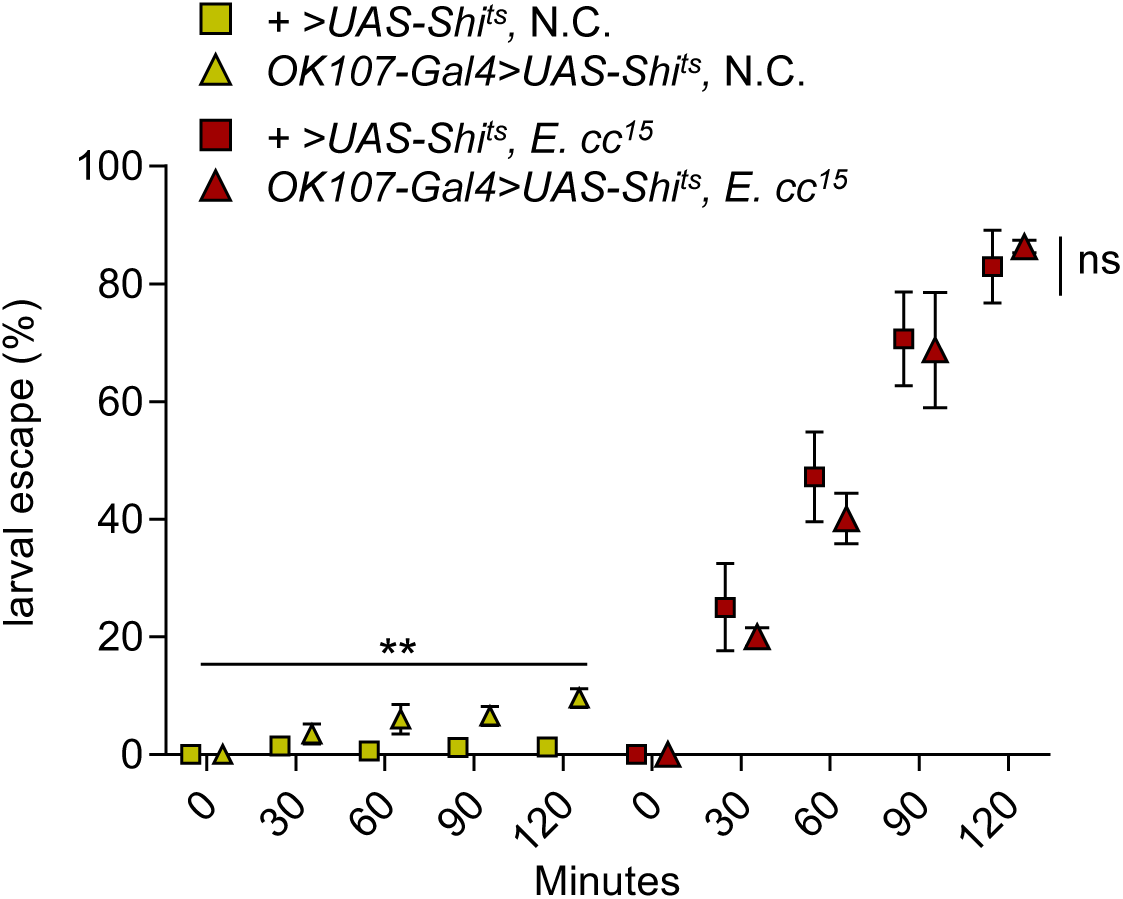
Inactivation of mushroom body intrinsic neurons does not affect larval escape from *E. cc^15^*-contaminated food. Escape behavior of control *(+>UAS-Shi^ts^*) and *OK107-GAL4>UAS-Shi^ts^* larvae exposed to uncontaminated (N.C.) or *E. cc^15^*-contaminated yeast at the restrictive temperature (31 °C). Statistical analyses were performed using a mixed-effects model (REML) with time and condition as fixed factors (ns = not significant; *p < 0.05; **p < 0.01; ***p < 0.001; ****p < 0.0001). Pairwise comparisons between selected conditions are shown.

**S10 Fig.**
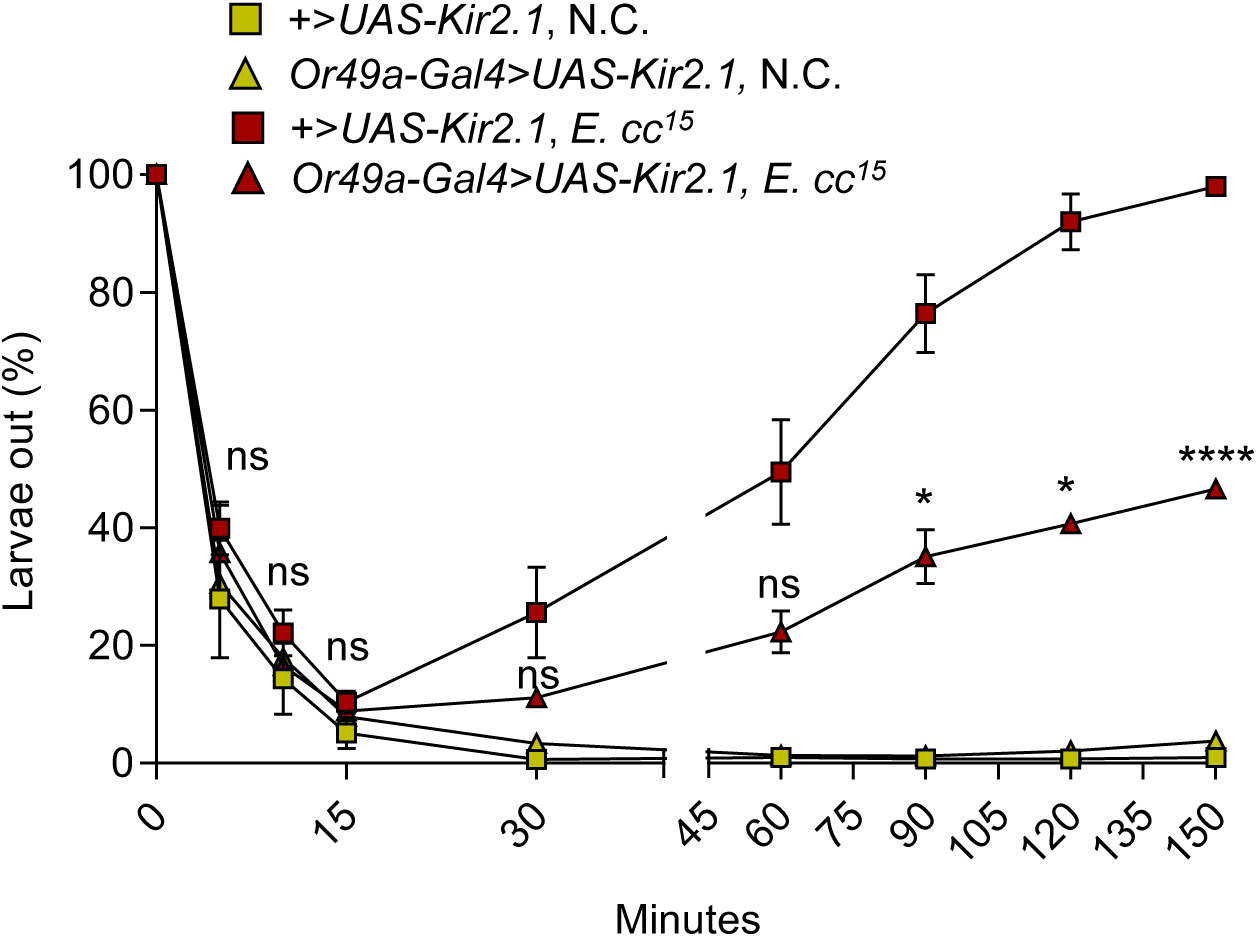
Silencing Or49a-positive neurons reduces *E. cc^15^*-induced dispersal without affecting locomotion or attraction to nutritive cues. Quantification of the proportion of larvae located outside the food source over time in control larvae (*+>UAS-Kir2.1*) and those in which *Or49a-*expressing neurons were silenced (*Or49a-GAL4>UAS-Kir2.1*). Larvae were placed at a fixed distance from either uncontaminated yeast or yeast contaminated with *E. cc^15^*. Statistical analyses were performed using a mixed-effects model (REML) with time and condition as fixed factors, followed by Tukey’s post-hoc tests (ns = not significant; *p < 0.05; **p < 0.01; ***p < 0.001; ****p < 0.0001). Pairwise comparisons between larvae *+>UAS-Kir2.1* and *Or49a-GAL4>UAS-Kir2.1* exposed to food contaminated with *E. cc^15^*are presented.

**S11 Fig.**
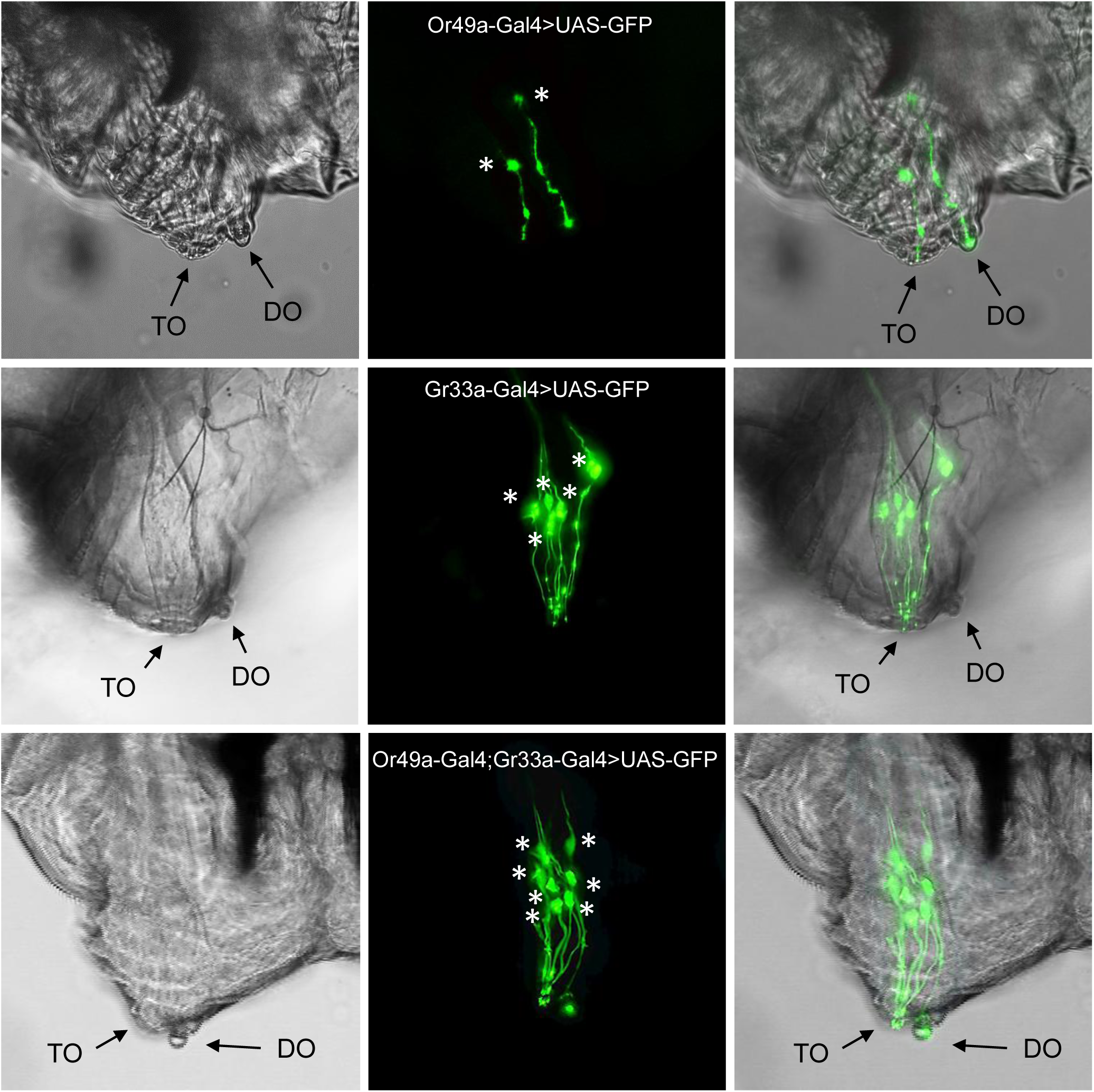
Determination of *Gr33a* and *Or49a* co-expression by double-driver analysis. Confocal images showing, from left to right, bright-field views of the dorsal and terminal organs (DO and TO; arrows), GFP expression driven by the indicated GAL4 lines, and merged images. Top: *Or49a-GAL4>UAS-6xGFP*. Middle: *Gr33a-GAL4>UAS-6xGFP*. Bottom: *Or49a-GAL4; Gr33a-GAL4>UAS-6xGFP*. Asterisks indicate neuronal cell bodies.

**S12 Fig.**
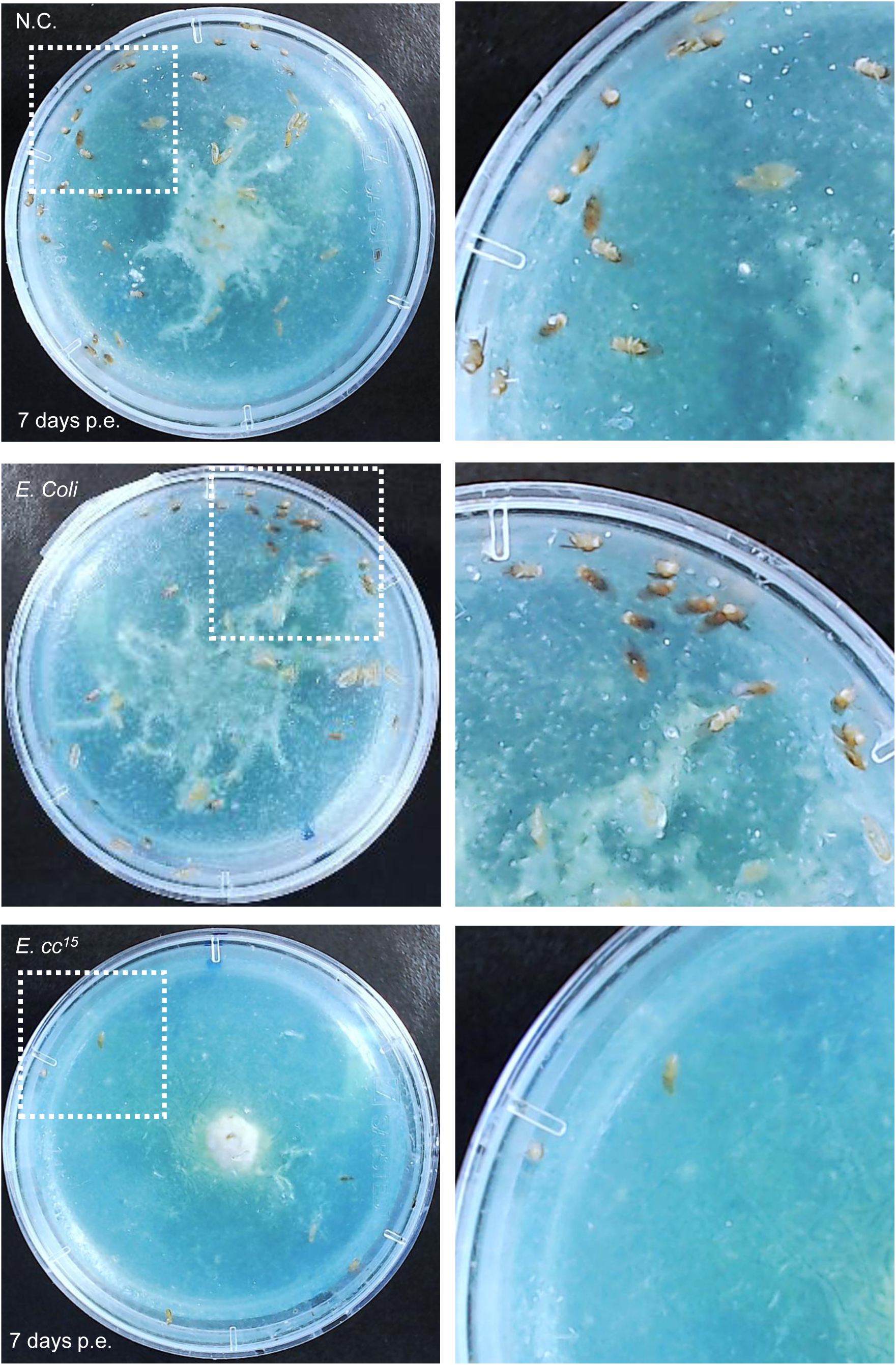
Larval exposure to *E. cc^15^*-contaminated food impairs development. Representative images showing adult emergence seven days after second-instar larvae were exposed to either uncontaminated yeast (N.C., top), yeast contaminated with *E. coli* (middle), or yeast contaminated with *E. cc^15^*(bottom), each applied at identical bacterial densities (OD = 40). Boxed regions are shown at higher magnification on the right. Images were acquired using a standard webcam

**S13 Fig.**
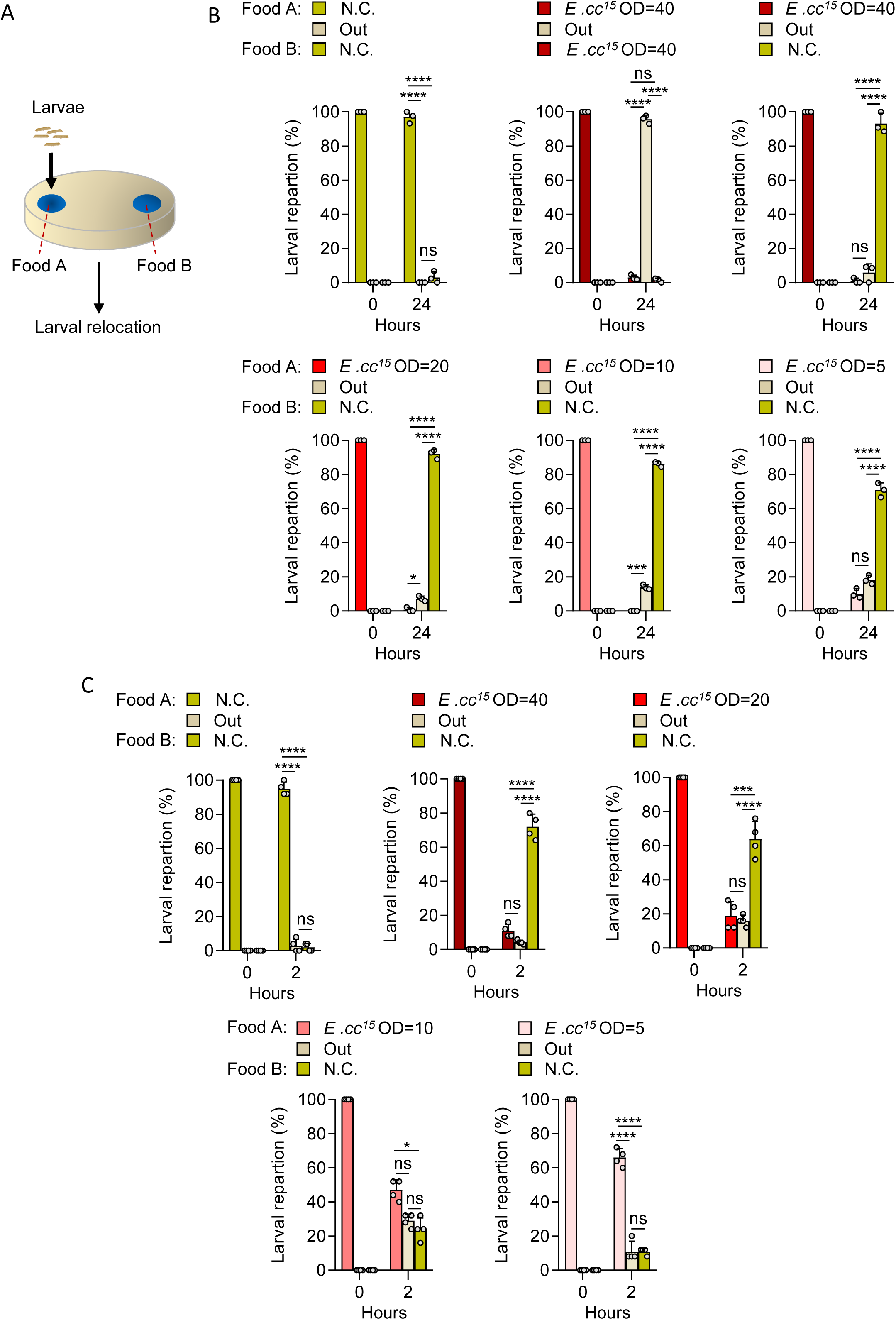
Larvae exposed to *E. cc^15^*-contaminated food relocate toward an uncontaminated food source. **(A)** Schematic overview of the assay used to monitor larval movement from an *E. cc^15^*–yeast mixture toward an uncontaminated yeast source. (B–C) Quantification of the proportion of larvae located on yeast contaminated with increasing concentrations of *E. cc^15^* (OD_600_ = 5, 10, 20, and 40) (Food A), in the surrounding agar region (Out), or on the uncontaminated yeast (Food B). Measurements were taken at 0 h and 24 h (B) or at 0 h and 2 h (C) following initial exposure. Both panels include a control condition in which Food A and Food B consisted of uncontaminated yeast, providing a baseline distribution in the absence of bacterial cues. Panel B additionally includes a second control in which both food patches were contaminated with *E. cc^15^* (OD_600_ = 40), establishing a baseline distribution when no clean alternative is available. Data are presented as means ± SD. Individual points represent experimental replicates (each consisting of groups of ≥ 25 larvae). Statistical analyses were performed using Fisher’s exact test (ns = not significant; *p < 0.05; **p < 0.01; ***p < 0.001; ****p < 0.0001).

**S14 Fig.**
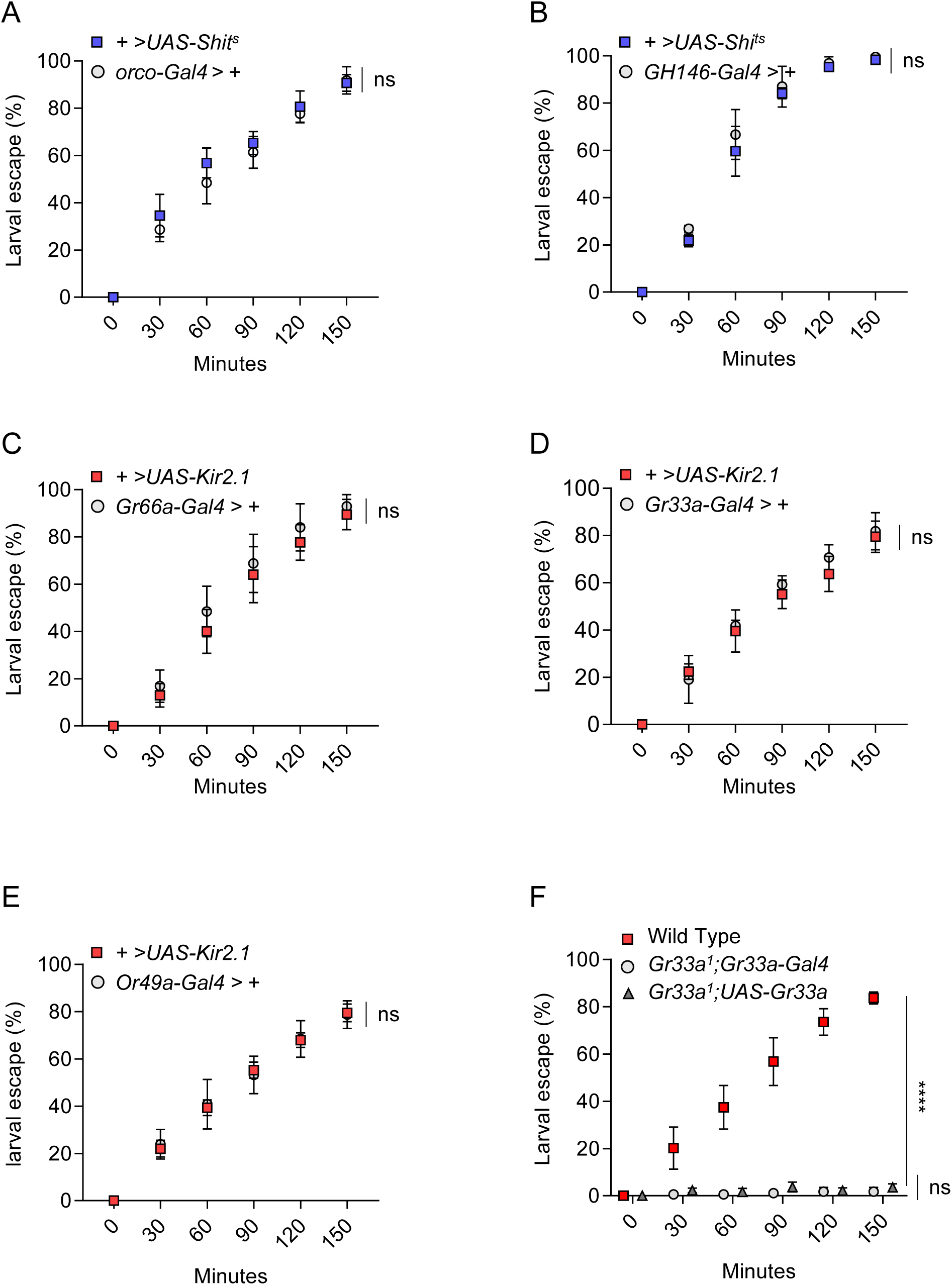
Baseline behavioral characterization of parental GAL4 and UAS lines on *E. cc^15^*-contaminated food. **(A-F)** Quantification over time of larval dispersal on *E. cc^15^*-contaminated yeast for parental lines used in the study. Panels show disperal of *+>UAS-Shi^ts^* and *Orco-GAL4* (A) or *GH146-GAL4* (B), *+>UAS-Kir2.1* controls and the corresponding parental GAL4 lines: *Gr66a-GAL4>+* (C), *Gr33a-GAL4>+* (D), or *Or49a-GAL4>+* (E), and wild-type, *Gr33a¹; Gr33a-GAL4* and *Gr33a¹; UAS-Gr33a* (F). Statistical analyses were performed using a mixed-effects model (REML) with time and condition as fixed factors (ns = not significant; ****p < 0.0001). Pairwise comparisons are shown.

---

**S1 Movie. Larval behavior on uncontaminated yeast.** Second-instar *Drosophila melanogaster* larvae (w¹¹¹^8^) were placed directly onto an uncontaminated yeast substrate. The video shows their behavior over the 3 h 30 min recording period. Images were acquired by time-lapse imaging (1 frame every 2 s) and compiled at 24 fps. https://figshare.com/s/54cf4674f0c1365e5b07

**S2 Movie. Larval behavior on *Ecc^15^*-contaminated yeast.** Second-instar larvae (w¹¹¹^8^) were placed directly onto yeast inoculated with *E. cc^15^* (OD_600_ = 40). The video shows their behavior over the 3 h 30 min recording period. Images were acquired by time-lapse imaging (1 frame every 2 s) and compiled at 24 fps. https://figshare.com/s/b143b90527a620300203

**S3 Movie. Larval behavior on *E. coli*-contaminated yeast.** Second-instar larvae (w¹¹¹^8^) were placed directly onto yeast inoculated with *E. coli* (OD_600_ = 40). The video shows their behavior over the 3 h 30 min recording period. Images were acquired by time-lapse imaging (1 frame every 2 s) and compiled at 24 fps. https://figshare.com/s/ebfbee471d060462bf31

## Notes

### Competing Interest Statement

The authors have declared no competing interest.

